# Emergence of Direction Selectivity at the Convergence of Thalamo-Cortical Synapses in Visual Cortex

**DOI:** 10.1101/244293

**Authors:** Anthony D Lien, Massimo Scanziani

## Abstract

Detecting the direction of an object’s motion is essential for our representation of the visual environment. Visual cortex is one of the main stages in the mammalian nervous system where motion direction may be computed *de novo.* Experiments and theories indicate that cortical neurons respond selectively to motion direction by combining inputs that provide information about distinct spatial locations with distinct time-delays. Despite the importance of this spatiotemporal offset for direction selectivity its origin and cellular mechanisms are not fully understood. We show that ~80+/−10 thalamic neurons responding with distinct time-courses to stimuli in distinct locations contribute to the excitation of mouse visual cortical neurons during visual stimulation. Integration of thalamic inputs with the appropriate spatiotemporal offset provides cortical neurons with the primordial bias for direction selectivity. These data show how cortical neurons selectively combine the spatiotemporal response diversity of thalamic neurons to extract fundamental features of the visual world.

---

The ability to detect the direction of motion of objects in the visual world is an essential property of sensory processing. In the primary visual cortex of mammals many neurons preferentially respond to stimuli moving in a specific direction within their receptive field ^1,2^. In primates and carnivores, this sensitivity to motion direction is believed not to be inherited from earlier stages of visual processing but computed *de novo* in visual cortex. Here, neurons likely extract directional motion from the visual scene by combining inputs that respond with different temporal delays to stimuli presented in different locations of the neuron’s receptive field, that is by combining spatially and temporally offset inputs ^3–11^. All theoretical models of direction selectivity, from the most general ^12,13^ to those that more closely mimic the receptive field structure of visual cortical neurons ^14–17^, are based on this spatiotemporal offset. However, the cellular mechanisms and synaptic connectivity patterns that generate the spatiotemporal offset of visual responses in cortical neurons remain speculative. Some models propose that direction selectivity emerges through intracortical interactions while others posit that direction selectivity is first generated through thalamo-cortical interactions. Other models, based on the rodents’ visual system, suggest that direction selectivity may actually not be computed in cortex but is inherited from earlier stages of visual processing. Models of direction selectivity generated through intracortical interactions (Fig. 1a) propose that anisotropic connectivity patterns between cortical neurons result in a spatial offset between excitation and inhibition or in a spatial offset between excitatory inputs with distinct time-courses ^10,15,16,18–22^. Models proposing that direction selectivity results from thalamo-cortical interactions suggest that spatially offset thalamic inputs with different time-courses in their responses to visual stimuli converge onto individual cortical neurons (Fig. 1a) ^7,23–26^. Models proposing that direction selectivity is not computed in the cortex, suggest that direction selectivity already existing in the retina may be transmitted to the cortex via the thalamus (Fig. 1a) ^27–30^. Here we take advantage of our ability to isolate individual thalamic inputs onto layer 4 (L4) cortical neurons of mice to identify the mechanism for the *de novo* generation of direction selectivity in primary visual cortex (V1).

**Figure 1.**
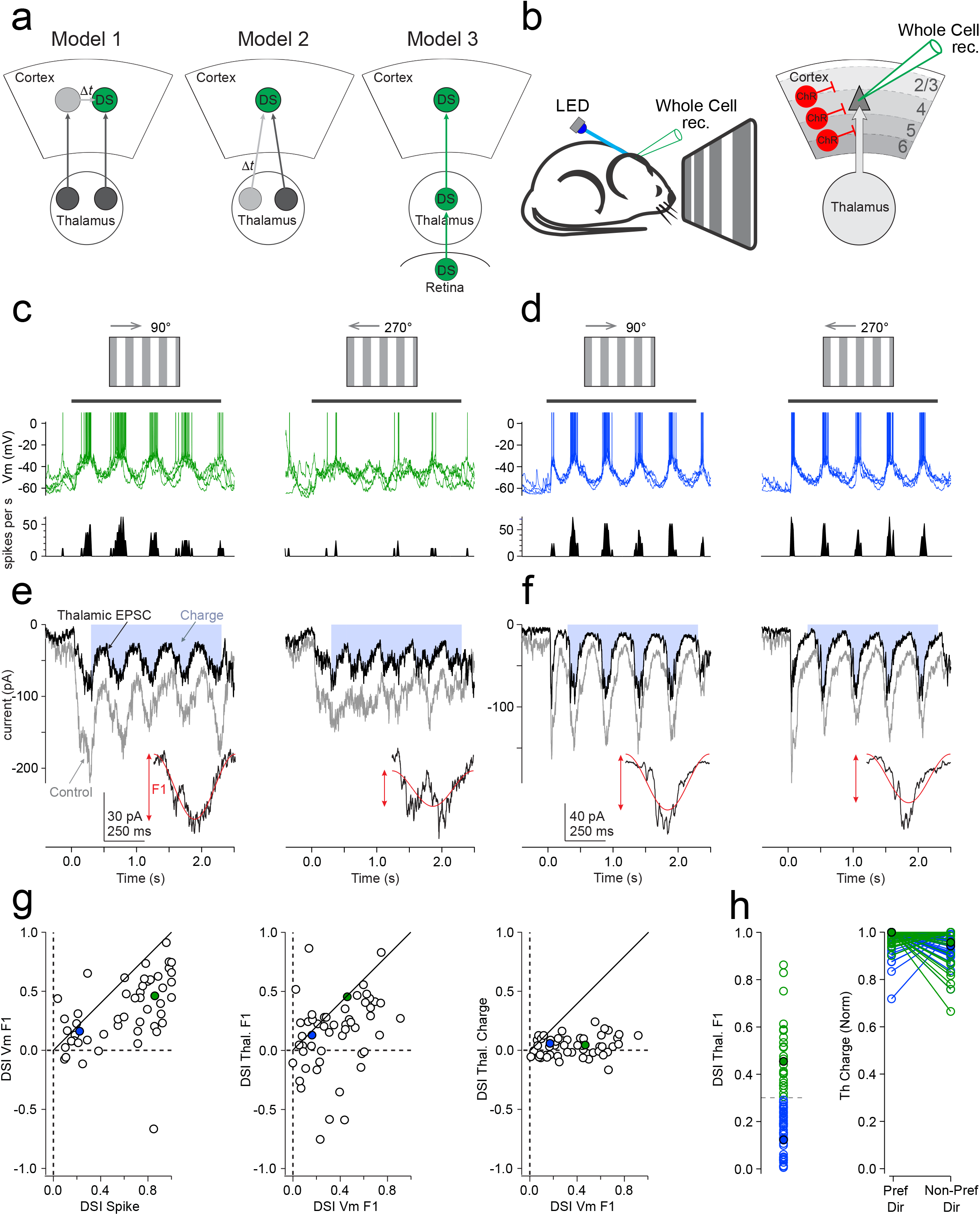
Amplitude modulation of thalamic excitation is direction selective. a. Two alternative models for the emergence of direction selectivity. Left: Model 1: The spatiotemporal offset of synaptic inputs onto a direction selective cortical cell (DS) is generated by intra-cortical circuits. Right: Model 2: The spatiotemporal offset is generated by the convergence of thalamic inputs with distinct spatial and temporal properties. b. Experimental configuration to isolate thalamic excitation: whole cell recordings in layer 4 of anesthetized mice during visual stimulation while photo-activating Channelrhodopsin expressing inhibitory neurons to silence cortex. c. Top: Current clamp recordings of a direction selective neuron in response to gratings drifting in the preferred (left) and non-preferred (right) direction (4 superimposed sweeps). Vm: membrane potential. Bottom: Peri-stimulus time histogram. d. As in (c) but for a non-direction selective neuron. e. Voltage clamp recording (V holding: -70 mV) of the neuron illustrated in (c) under control conditions (gray traces) and during cortical silencing (black traces) to isolate thalamic excitation. Inset is a cycle average and the red trace a sinusoidal fit. The arrow (F1) shows the amplitude of the modulation. The blue shaded area is the current integral used to estimate the thalamic excitatory charge. f. As (e) but for neuron illustrated in (d). g. Summary plots for 52 recordings (31 mice). The scatter plots show the relationship between the direction selectivity index (DSI) of the spikes (DSI spike), the F1 modulation of the membrane potential (DSI Vm F1), the F1 modulation of thalamic excitation (DSI Thal. F1), and the thalamic charge (DSI Thal. Charge). Negative values on the y-axis illustrate opposite direction preference as compared to index of x-axis. Note the absence of correlation between the DSI of the thalamic charge and the F1 modulation of Vm. Green and blue filled data points refer to the neurons in (c,e) and (d,f), respectively. h. Left: Distribution of DSIs of the F1 modulation of thalamic excitation for all recorded neurons. 66 recordings (41 mice). Above a DSI of 0.3 (horizontal dotted line) the thalamic excitation is considered direction selective. Right: The charge of thalamic excitation in the preferred and non-preferred directions for each recorded neuron. For each cell the charge is normalized by the largest charge. Filled data points are neurons from (c,e) and (d,f).

## Thalamic excitation reports direction of motion

To investigate the synaptic mechanisms underlying direction selectivity, we performed whole-cell patch-clamp recordings from neurons located between 300-550 um from the pial surface in V1 of anesthetized mice while displaying visual stimuli to the contralateral eye ^31^. This range of depth corresponds largely to the radial extent of L4 but may include some deep L2/3 and superficial L5 neurons. For simplicity we will refer to the neurons recorded at this depth as L4 neurons. We recorded the spiking and the membrane potential (Vm) in response to 12 different stimuli presented on a monitor and consisting of gratings (arrays of dark and light bars) of 6 orientations drifting in either one of two opposite directions along the axis perpendicular to the gratings’ bars. When presented with gratings at their preferred orientation some L4 neurons (Fig. 1c) fired many more action potentials in response to movement in one direction as compared to the opposite, consistent with previous studies ^32^. Vm fluctuated at the fundamental temporal frequency of the drifting grating (F1 modulation), that is at the frequency at which any pixel on the monitor transitions from dark to light and back to dark (Fig. 1c and d). The direction selectivity index (DSI, see methods) of the amplitude of the F1 modulation of Vm (VmF1) correlated with the DSI of the neuron’s spiking response (r=0.52 p=0.000103, Fig. 1g, left) and in most cells the preferred direction was the same for both parameters (46/51 matching preferred direction p<0.0001 binomial test; Fig. 1g left).

To determine whether the direction preference of L4 neurons is already apparent in the thalamic excitatory synaptic input we isolated thalamic excitation by silencing cortical excitatory neurons via photo-activation of cortical inhibitory interneurons expressing Channelrhodopsin 2 (ChR2) ^33^, while recording from L4 neurons in the voltage clamp configuration at the reversal potential of inhibition, as described previously ^31^ and presenting gratings drifting in either of the two direction perpendicular to the previously determined preferred orientation. Cortical silencing completely abolished firing in cortical excitatory neurons, including in the deepest cortical layers, and reduced visually evoked excitation by ~65% (63±16%, drifting grating; n=66 cells; 68±16%, static grating; n=53 cells; Extended Data Fig. 1), consistent with the fact that during visual stimulation the majority of excitation received by L4 neurons is of cortical origin^26,31,34^. The residual excitation represents the isolated thalamic input^31,34^.

To determine whether thalamic excitation shows direction selectivity we used two parameters: First, the amplitude of the F1 modulation of the thalamic excitatory current. Second, the thalamic excitatory current integrated over the duration of the visual stimulus (thalamic charge), i.e. the total amount of thalamic excitation received by a L4 neuron in response to the stimulus. The amplitude of the F1 modulation of thalamic excitation (ThalF1) showed direction selectivity in 38% of the neurons (we defined ThalF1 as being direction selective when the DSI exceeded 0.3; 25/66 cells; Fig. 1h left). Importantly, the preferred direction of ThalF1 predicted the preferred direction of VmF1 in the majority of cells (37/52 cells p=0.0032; binomial test; Fig. 1g middle; if the test is restricted to cells with DSI VmF1>0.3: 22/28 had the same preferred direction; p=0.0037, binomial test; the DSI of ThalF1 also correlated with the DSI of VmF1 [r=0.33; p=0.0168; n=52 cells; Fig. 1g middle]).

In contrast, the thalamic charge showed no direction selectivity (in 64/66 cells DSI did not exceed 0.3) and was on average the same for the preferred and non-preferred directions of ThalF1 (for cells with DSI ThalF1 > 0.3 the normalized thalamic charge was 0.98 ± 0.03 in the preferred and 0.93 ± 0.09 in the non-preferred direction, p=0.13 n=25 cells; for DSI ThalF1 < 0.3: 0.97 ± 0.06 preferred, 0.95 ± 0.06 non-preferred, p = 0.22 n=41 cells, Wilcoxon rank-sum test, Fig. 1h right). The DSI of thalamic charge did not correlate with the DSI of VmF1 (r=0.2 p=0.16 n=52 cells, Fig. 1g right) and did not predict the preferred direction of VmF1 (32/52 had the same preferred direction; p: 0.126, binomial test; if the test is restricted to cells with DSI VmF1>0.3: 18/28 had the same preferred direction; p=0.1849, binomial test; Fig. 1g right). Finally, to quantify the possible contribution of a thalamic charge bias to the DSI of ThalF1 we equalized the thalamic charge evoked by gratings drifting in both directions (see methods). This procedure led to a very small, yet significant reduction in DSI of ThalF1 (subset of neurons where DSI of ThalF1 > 0.3: DSI before equalization: 0.49 +/− 0.15; DSI after equalization: 0.46 +/− 0.17; p=0.022, paired t-test; n=25; Extended Data Figure 2). These data demonstrate that direction preference in L4 neurons is already prominent in the amplitude modulation of thalamic excitation ^26^ but much less so in the charge. Thus, an essentially equal quantity of thalamic excitation is differently distributed in time depending on the direction of the stimulus.

## The spatiotemporal offset of thalamic excitation

What accounts for the differential temporal distribution of thalamic excitation for stimuli moving in opposite directions? The amplitude of thalamic excitation in response to a moving stimulus is the summation of the momentary thalamic excitation driven by the current position of the stimulus with the residual excitation generated by the previous stimulus position within the receptive field. Heterogeneity in the time-course of thalamic excitation elicited by stimuli at different receptive field locations could cause the observed differences in the amplitude modulation for motion in opposite directions ^7,23–26^.

To determine whether the time-course of thalamic excitation depends on the position of the stimulus, we presented, while silencing cortex, static gratings of the preferred orientation at 16 different spatial phases, separated by 22.5° and presented randomly (250 ms each; Fig. 2a-b). The static gratings triggered thalamic excitatory postsynaptic currents (EPSCs) with a latency of 39.8 ± 7.3 ms (time to 20% of peak amplitude, n=53 neurons), an amplitude of -105 ± 61 pA (range -28 to -280 pA, measured for the spatial phase eliciting the largest amplitude, n=53 neurons) and a duration of 145± 33 ms (range: 46-180 ms duration; n=53 neurons; time from 10% to 90% of integral).

**Figure 2.**
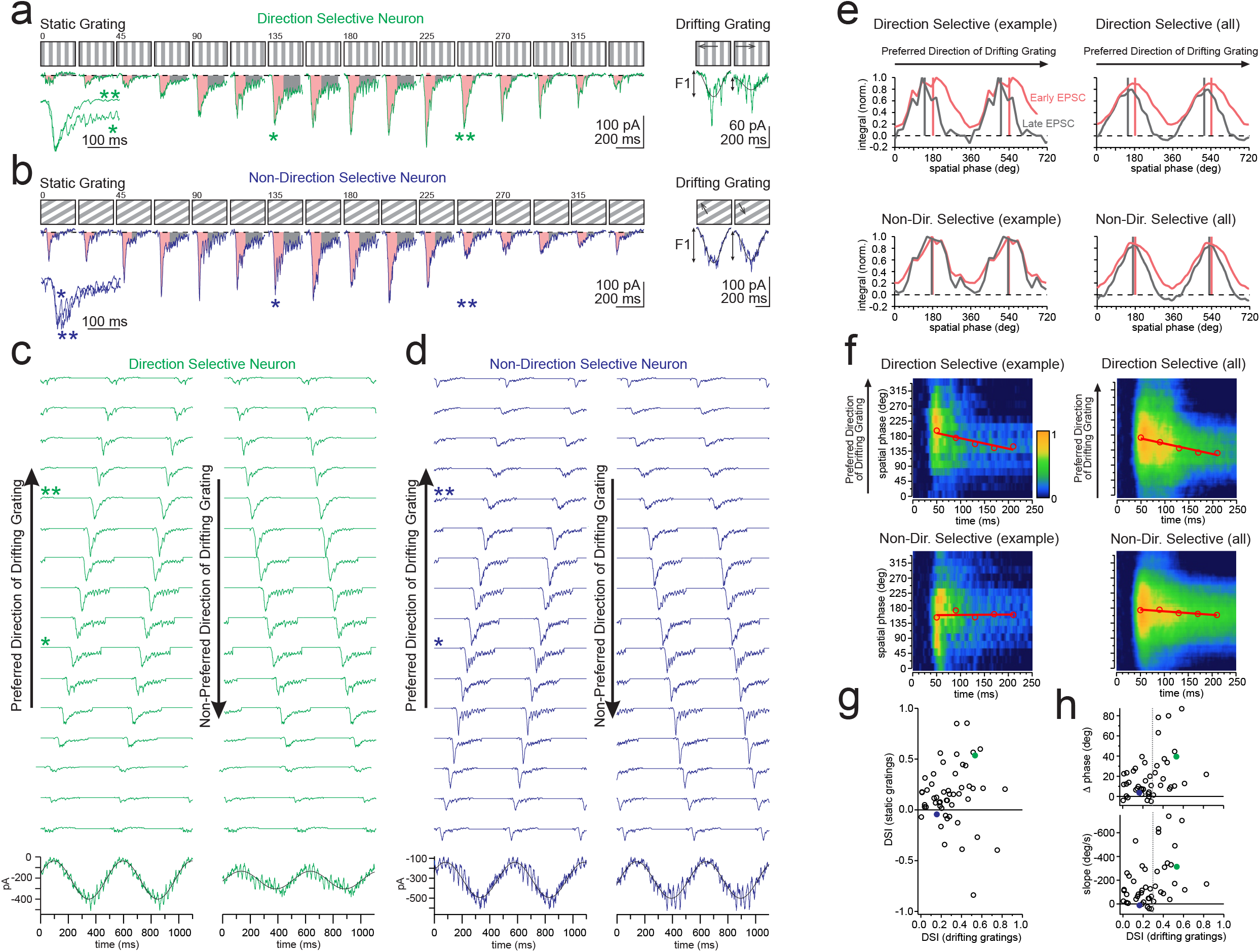
Time-course of thalamic excitation to static stimuli explains direction selectivity to moving stimuli. a. Thalamic EPSCs recorded in a L4 direction selective neuron (DSI F1 thalamic excitation: 0.53) in response to static gratings presented at 16 different phases (0-337.5°, 22.5° spacing) of the preferred orientation (250 ms duration). The early (30-100 ms after stimulus onset) and late (110-230 ms after stimulus onset) portion of the EPSC are shaded in pink and gray, respectively. The right inset is a cycle average of thalamic excitation recorded in the same neuron in response to a drifting grating in the preferred (left) and non-preferred (right) direction and the black trace a sinusoidal fit. The arrow (F1) shows the amplitude of the modulation. The left inset shows, on an expanded time-scale the superimposition of two example thalamic EPSCs (asterisks; peak scaled) elicited by static gratings of two different phases. Note the different decay time-course. b. As in (a) but for a non-direction selective neuron (DSI F1 thalamic excitation: 0.16). c. Each green trace is the thalamic EPSC to one of the 16 phases of the static grating of the neuron in (a). Each trace is the concatenation of identical EPSCs separated by 500 ms, i.e. a full cycle of the drifting grating. Traces are staggered in time relative to one another by 31.25 ms, i.e. a 16^th^ of a full cycle of the drifting grating. Asterisks mark EPSC shown in (a). Traces on the left are ordered according to the sequence of phases of a grating drifting in the preferred direction. Traces on the right are ordered according to the non-preferred direction. The bottom trace is the algebraic sum of the green traces above and the black trace is a sinusoidal fit. Note the pronounced F1 modulation of the left as compared to the right trace. d. As in (c) but for the non-direction selective neuron in shown in (b). e. Phase dependent modulation of the integral of the early (pink) and late (gray) portion of the thalamic EPSC. Top left: example neuron in (a). Bottom left example neuron in (b). The cycle is repeated twice for clarity. Right: summary for all neurons with DSI> 0.3 (top; n = 18) and DSI<0.3 (bottom; n = 28). Plots were aligned so that the preferred spatial phase of the early EPSC (estimated by vector averaging, see methods) occurred at 180° and ordered such that increasing spatial phase corresponded to the drifting grating preferred direction. Example neurons from (a) and (b) were shifted by +7.2° and -18.75° degrees, respectively. Note the phase shift between the early and late portion of thalamic EPSCs for neurons with DSI>0.3 (vertical bars aligned at the peak of the cycle). f. Spatiotemporal receptive field of thalamic EPSCs to static gratings. Hotter colors signify larger amplitudes (see scale to the right). Red circles mark the phase with the largest excitation at a given time. The red line is a linear fit. Top left: example neuron in (a). Bottom left example neuron in (b). Right: summary for all neurons with DSI> 0.3 (top; n = 18) and DSI<0.3 (bottom; n = 28). Population heat-maps were aligned so that the preferred spatial phase of the earliest time bin occurred at 180° and ordered such that increasing spatial phase corresponded to the drifting grating preferred direction. Note the different slopes of the linear fit. g. The DSI in response to drifting gratings is plotted against the DSI computed by the summed responses to static gratings. 53 recordings (34 mice). Note that for most experiments the direction preference to drifting gratings matches the direction preference of the response computed by summing the response to static gratings. Filled circles corresponds to cells in (c) and (d). h. Top: The phase difference between the early and late portion of the thalamic EPSC (see (e)) is plotted against the DSI in response to drifting gratings. Bottom: The slope of the linear fits (see (g)) is plotted against the DSI in response to drifting gratings. 46 recordings (32 mice). The vertical dotted line marks DSI=0.3.

To validate that static gratings allow us to recapitulate the dynamics of thalamic excitation in response to drifting gratings we computed the algebraic sum of thalamic EPSCs evoked by each of the 16 phases of the static gratings (Fig. 2c,d). The currents were staggered in time (by 31.25 ms, i.e. by 1/16 of the period of the drifting grating) to match the temporal sequence at which each of the 16 individual spatial phases occur during a drifting grating (Fig. 2c,d); we also staggered the gratings by shorter intervals to mimic higher temporal frequencies; Extended Data Fig. 3). We compared the algebraic sum in which EPSCs were ordered according to the spatial phase sequence simulating the motion of the grating in one direction against the sum simulating motion in the opposite direction. In the large majority of recorded L4 neurons the F1 amplitude modulation of the summed thalamic EPSCs evoked by static gratings had a direction preference matching that of ThalF1 evoked by drifting gratings (Fig. 2g; 41/53 matching preferred direction p<0.0001, binomial test; for neurons where DSI ThalF1 to drifting gratings >0.3: 18/23 matching preferred direction p=0.01; binomial test; among the 12 neurons where direction preference was not matched 7 were not direction selective (DSI ThalF1 to drifting gratings <0.3)). While we have no explanation for the unmatched directional preference of the remaining 5 cells, the thalamic charge of these neurons in response to drifting gratings was on average not direction selective (DSI of thalamic charge: 0.09 +/− 0.07; n = 5) consistent with the data above). These results validate static gratings as an approach to determine the dynamics of thalamic excitation underlying direction preference. Below we restrict our analysis to those 46 neurons where the DSI to static gratings is > -0. 1 (Fig. 2g).

The amplitude of thalamic EPSCs evoked by static gratings depended on the spatial phase of the grating, consistent with the spatial separation in ON and OFF sub-regions of thalamic excitation ^31^ (Fig. 2a). Strikingly, however, the time-course of the decay of the thalamic EPSCs also depended on the spatial phase (Fig. 2a, asterisks). We quantified the impact of the spatial phase of the static grating on the time-course of the thalamic EPSC by measuring the integral of the early (30-110 ms from stimulus onset, Fig. 2a-b shaded pink regions) and late (110-230 ms, Fig. 2a-b shaded gray regions) portion of the EPSC, to capture the peak and the decay, respectively (Fig. 2e). Both the magnitude of the early and the late EPSC fluctuated sinusoidally with a period spanning the 16 phases of the static gratings (i.e. one whole cycle). Crucially, the phase relationship between the modulation of the magnitude of the early and late EPSC was not fixed but varied from cell to cell (Fig. 2h top), ranging from 0° difference (i.e. the spatial phase that triggers the largest early EPSC also triggers the largest late EPSC) to 87° difference (i.e. the spatial phase that triggers the largest early EPSC is shifted relative to the phase that triggers the largest late EPSC). We discovered that the phase relationship between the early and late thalamic EPSC predicted the direction preference of the cell to drifting gratings. The predicted preferred direction was the direction of movement in which the spatial phase eliciting the largest late EPSC preceded the spatial phase eliciting the largest early EPSC. Intuitively, two consecutive EPSCs with distinct decays will summate to produce a larger peak current if the slow one precedes the fast one than if the fast one precedes the slow one. In 89% of the neurons (41 out of 46), the predicted preferred direction matched that observed with drifting gratings (p < 0.0001; Binomial test). In those few cells where the phase relationship did not predict the preferred direction of drifting gratings, the phase difference was close to 0° and the cells were poorly direction selective (DSI ThalF1 < 0.3; the phase difference between the early and late EPSCs correlated with the DSI in response to drifting gratings [r=0.405 p=0.0053; n=46 cells; phase difference for DSI ThalF1 < 0.3: 10.8 ± 11.5°; n=28 cells; phase difference for DSI ThalF1 > 0.3: 33.8 ± 26.9°; n=18 cells; p = 0.000603, Wilcoxon rank-sum test] Fig. 2h, top). To visualize the phase dependent shift in the early and late EPSC we computed the spatiotemporal receptive field that is, a heat map in which the time-course of the thalamic EPSC is plotted for each spatial phase of the static grating (Fig. 2f). For each time bin, we identified the phase with the largest excitation and fitted a linear function through the data. Steeper slopes indicate larger phase differences between the early and late EPSC. Negative slopes predict that the preferred direction is for gratings drifting in the direction of increasing spatial phase. Again, in 89% of the neurons (41 out of 46), the predicted preferred direction matched that observed with drifting gratings (p < 0.0001; Binomial test; the slope correlated with the DSI in response to drifting gratings [r = -0.38; p=0.009, n=46 cells; slope for DSI ThalF1 < 0.3: -104 ± 129°/s; n=28 cells phase difference for DSI ThalF1 > 0.3: -311 ± 232°/s; n=18 cells p = 0.00204 Wilcoxon rank-sum test] Fig. 2h, bottom). These results demonstrate that different positions of the stimulus trigger thalamic excitation with different early and late components. Importantly, the difference between the spatial positions triggering the largest early and the largest late component provides the initial bias for a preferred direction of motion in L4 neurons.

## Combining individual thalamic inputs with distinct spatiotemporal response properties

By what mechanism is the time-course of thalamic excitation modulated by the spatial phase of the grating? We tested the possibility that the time-course of thalamic excitation depends on the time-course of the firing of the thalamic neurons they receive input from. For this we needed to record from synaptically connected thalamo-cortical pairs ^35,36^ and compare the time-course of firing of presynaptic thalamic neurons with the time-course of decay of thalamic excitation. We performed extracellular recordings from thalamic neurons using four shank linear probes inserted in the dorsolateral geniculate nucleus (dLGN), the primary thalamic relay from the retina to V1, while simultaneously recording thalamic excitation from L4 neurons during cortical silencing. The response of thalamic neurons to static gratings varied from transient to sustained, consistent with previous reports^37^ (Extended Data Fig. 4). While the firing rate of thalamic neurons was modulated by the phase of the grating, the type of response, i.e. transient or sustained, remained the same irrespective of phase (Extended Data Fig. 4).

Monosynaptic connections between thalamic neurons and L4 cortical neurons were identified based on latency, time course, and probability of events in the cortical neuron following spikes in individual thalamic neurons during the presentation of drifting gratings under cortical silencing conditions (see methods and Extended Data Fig. 5). Out of 40 L4 whole cell recordings in 24 mice and a total of 739 isolated thalamic units, 23 thalamo-cortical pairs, recorded in 15 mice satisfied those criteria and were considered monosynaptically connected. Those 23 connected pairs consisted of 17 L4 neurons and 23 thalamic units, because while for most L4 neurons we found only one presynaptic thalamic unit, for two L4 neurons we isolated two presynaptic units and in two L4 neuron we isolated three presynaptic units (Extended Data Fig. 6). Unitary EPSCs (uEPSCs) had an average latency of 2.09 ± 0.51 ms (peak time of the peri-spike time histogram (PSpTH, see methods and Extended Data Fig. 5)), an average and median peak amplitude of -10.8 ± 11.6 pA and 6.4 pA, respectively (range -1.8 to -48.4 pA), and a jitter of 211 ± 47μs (half-width at half-max of PSpTH; n=23 connected pairs; uEPSPs had a mean and median amplitude of 0.81+/−0.79 mV and 0.57 mV, respectively and ranged from 0.14 - 3.4 mV; see methods and Extended Data Fig. 5). These thalamo-cortical connected pairs allowed us to compare the time course of thalamic excitation recorded in the postsynaptic L4 neurons with the time course of firing of their presynaptic thalamic input neurons.

The time course of firing in presynaptic thalamic units matched the duration of thalamic excitation in postsynaptic L4 neurons. The example in Figure 3 shows two units, 1 and 2, isolated in the thalamus, both converging on the same cortical neuron recorded in L4 (Fig. 3a). These two units responded maximally to distinct phases of the static grating (Fig. 3c, bottom). Furthermore, these two units also had distinct response types, unit 1 showing transient and unit 2 sustained responses to the stimulus. Spatial phases of the static grating eliciting transient activity in unit 1 and little or no activity in unit 2 also elicited fast decaying thalamic excitation in the L4 neuron (Fig. 3d, left). In striking contrast, spatial phases of the grating that triggered sustained activity in unit 2 elicited slow decaying thalamic excitation in the L4 neuron (Fig. 3d, right). Thus, the time-course of firing of these two thalamic neurons matched the time-course of thalamic excitation in their postsynaptic L4 neuron.

**Figure 3.**
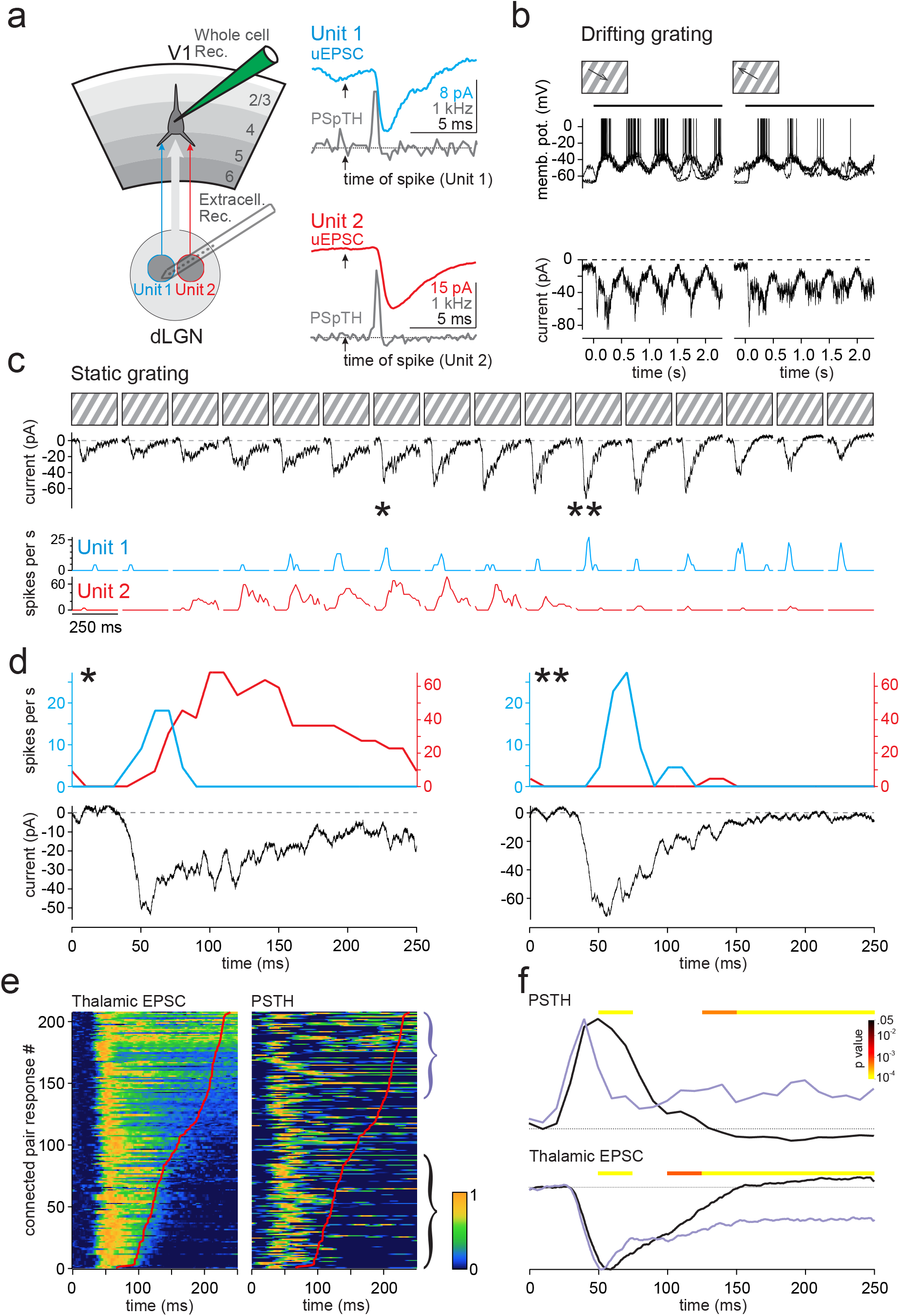
The time-course of firing of thalamic neurons explains the time-course of thalamic excitation. a. Left: Schematic of experimental configuration. Right: Two unitary excitatory postsynaptic currents (uEPSC; blue and red traces) from two presynaptic thalamic units (Unit 1 and 2) recorded in the same L4 cortical neuron while silencing cortex. The arrows indicate the time of the spike of the respective unit. The gray trace is the peri spike time histogram (PSpTH) of events recorded in the L4 neuron (see methods and Extended Data Fig. 5). The increase in the frequency of events shortly after the spike of the unit corresponds to the onset of the uEPSC. The dotted line marks an event frequency of 0.25 kHz. b. The direction selective response of the L4 neuron in (a) to gratings drifting in the preferred (left) and non-preferred (right) direction. Top: current clamp recordings (action potentials are truncated). Bottom: Voltage clamp recordings during cortical silencing to isolate thalamic excitation. c. Responses to static gratings presented at 16 different phases of the preferred orientation of the neuron in (c). Top row: Thalamic excitation recorded in the L4 neuron (same as (b)). Bottom row: Peri-stimulus time histograms (PSTHs) of the same two thalamic units as (a). d. Responses to two distinct phases of the static grating (from (c)) on expanded axes. Note that the phase of the static grating eliciting thalamic excitation with a slow decay (left) triggers a transient and a sustained response in units 1 and 2, respectively. The phase eliciting a fast decaying thalamic excitation (right) triggers a response in unit 1 only. e. Heat-maps of responses to static gratings for 20 thalamo-cortical connected pairs (15 postsynaptic L4 cortical neurons; 20 presynaptic thalamic units; 13 mice; only pairs where the thalamic unit responded to static gratings were included; see methods). Each row on the left panel is the amplitude of thalamic excitation recorded in one of the 15 L4 cortical neurons in response to one of the 16 phases of a static grating presented at the preferred orientation for that neuron. The rows are ordered according to the duration of thalamic excitation. The red line is the time at which the integral (charge) of the thalamic excitatory response reaches 90% of total. The amplitude of the PSTHs of the simultaneously recorded presynaptic thalamic units are shown on the corresponding rows on the right panel. Note that slower thalamic excitation (left panel, top) is accompanied by longer lasting PSTHs (right panel, top). The red line on the right panel is a copy of the red line on the left panel to facilitate comparison. Heatmap values are smoothed along the y-axis. f. Top: Average PSTHs of the top and the bottom rows of the right heat-map in (f) are shown in purple and black respectively (brackets in (e) illustrate the range of rows used for the average PSTHs). Bottom: average thalamic excitation for the same rows. P-values are for Wilcoxon rank-sum test comparing black and purple traces in 20ms bins.

To compare the time-course of thalamic excitation recorded in L4 neurons with the time-course of the firing of their presynaptic thalamic neurons across experiments we created a heat-map of the thalamic EPSC amplitudes ranked according to their duration (Fig. 3e, left; see Methods). Every row in the heat-map reports the amplitude of the thalamic EPSC in one L4 neuron in response to one of the 16 spatial phases of the grating. Every row in the adjacent heat-map (Fig. 3e, right) shows the amplitude of the simultaneously recorded peri-stimulus time histogram (PSTH) of the corresponding presynaptic thalamic neuron. There was a marked and highly significant correlation between the time-course of the thalamic EPSCs in L4 neurons and the time-course of the firing of their presynaptic thalamic neurons (average pairwise Pearsons correlation: 0.40; n=208 paired EPSC/PSTH spatial phase responses; significantly different than correlation of shuffled PSTHs; p<0.0001; see methods). Accordingly, the time-course of the average PSTH of thalamic neurons firing in response to static gratings triggering slow thalamic EPSCs was significantly more sustained than the time-course of the average PSTH in response to gratings eliciting fast EPSCs (Fig. 3f). Thus, the time-course of thalamic excitation onto a L4 neuron depends on the spatial phase and follows the time-course of the activity of its presynaptic thalamic neurons.

These results suggest that L4 neurons extract motion direction by combining thalamic inputs with distinct spatial and temporal response properties. In other words, the spatiotemporal receptive field of thalamic excitation of a L4 cortical neuron (e.g. Fig. 2h) should reflect the spatiotemporal activity pattern of the combined population of its presynaptic thalamic neurons. Consistent with this possibility even the combined spatiotemporal activity pattern of units 1 and 2 (from the example above) approximates the spatiotemporal receptive field of their common L4 target neuron (Fig. 4a), with the more sustained unit 2 being active at lower and the transient unit 1 being active at higher spatial phases. Because recordings of multiple thalamic units that are converging on a simultaneously recorded cortical neuron are rare, we verified the above hypothesis by generating a “compound”, direction selective cortical neuron and its set of thalamic inputs. We combined all eight thalamo-cortical connected pairs in which the cortical neuron had a DSI of thalamic excitation larger than 0.3 (total of six L4 neurons: four neurons with one presynaptic thalamic unit, two neurons with two presynaptic thalamic units; Fig. 4b). We aligned and averaged the spatiotemporal receptive fields of thalamic excitation of the six cortical neurons to obtain the spatiotemporal receptive field of our compound cortical neuron (Fig. 4b). The spatiotemporal receptive field of the compound neuron had a negative slope, consistent with the fact that the preferred directions of its six constituent neurons was for gratings drifting in the direction of increasing spatial phase. Strikingly, the resulting combined spatiotemporal activity pattern of the eight presynaptic thalamic inputs also had a negative slope: while the more sustained units were active at the lower spatial phases, the higher spatial phases were dominated by transient thalamic units. Thus the combined spatiotemporal activity pattern of the eight presynaptic thalamic units predicted the direction preference of the compound postsynaptic neuron (Fig. 4b).

**Figure 4.**
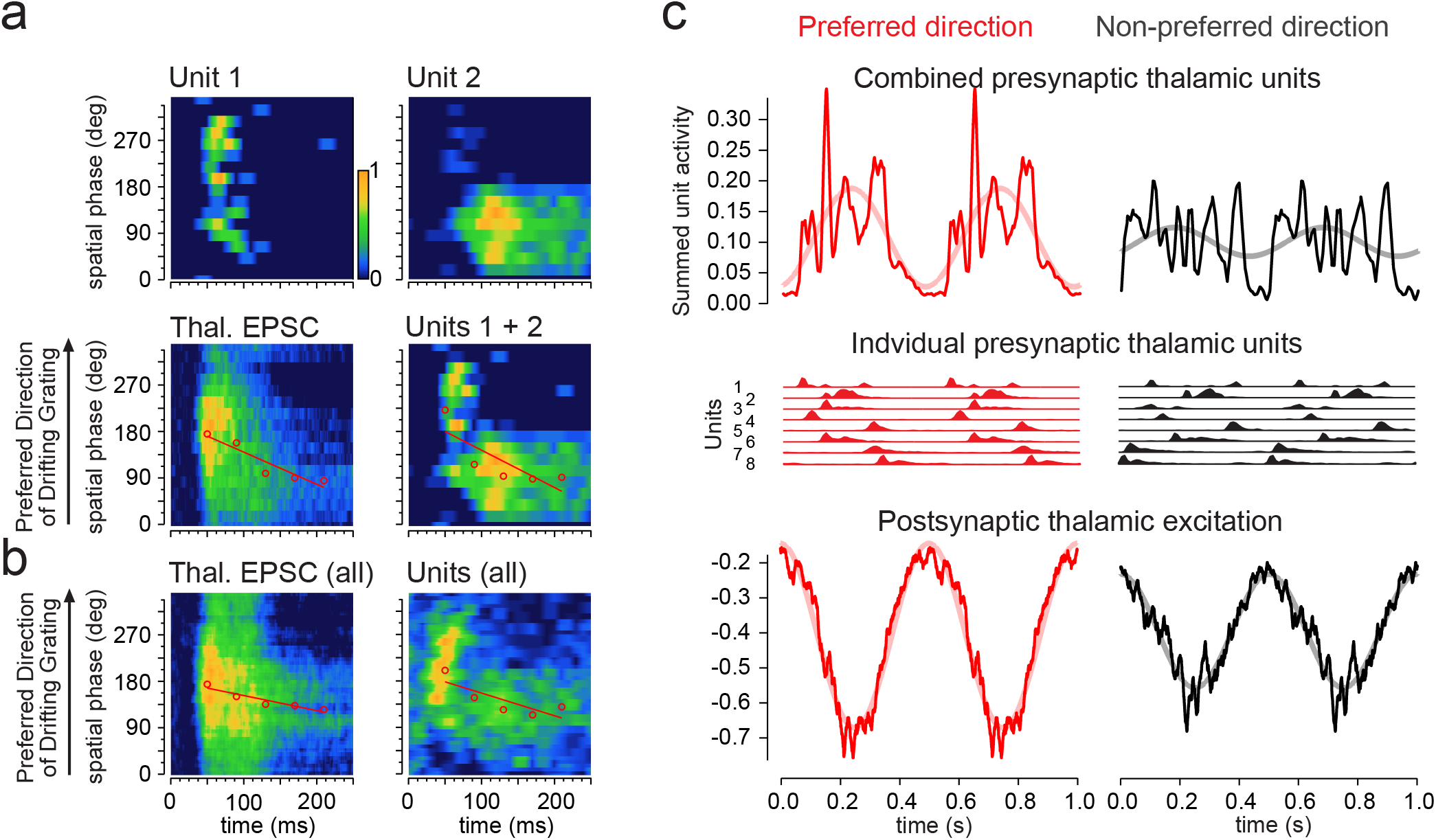
The temporal distribution of thalamic activity underlies cortical direction selectivity. a. Top: Spatiotemporal receptive field of the firing of thalamic unit 1 (left) and 2 (right; same units as in Fig. 3d) in response to static gratings. Note the phase difference and the different time-course in the responses between the two units. Bottom, left: Spatiotemporal receptive field of thalamic excitation recorded in a L4 cortical neuron (same neuron as in Fig. 3d) that was postsynaptic to units 1 and 2. Right: Combined spatiotemporal receptive field of units 1 and 2. Note the similarity between the summed spatiotemporal receptive field of the firing of units 1 and 2 and the spatiotemporal receptive field of thalamic excitation. b. Left: Average spatiotemporal receptive field of thalamic excitation to static gratings from 8 pairs where the cortical L4 neuron had a DSI > 0.3. Spatiotemporal receptive fields were aligned so that the preferred spatial phase of the earliest time bin occurred at 180° and ordered such that increasing spatial phase corresponded to the drifting grating preferred direction. Right: Combined spatiotemporal receptive field of the firing of the 8 thalamic units presynaptic to the 6 cortical neurons. c. Top: Summed activity of the same 8 presynaptic thalamic units to gratings drifting in the preferred (left, red traces) and non-preferred (right, black traces) direction (two identical cycles are shown for clarity; pink and gray lines are sinusoidal fits). The PSTHs of each of these units is shown below (units 1 and 2 correspond to units 1 and 2 in (a)). The units were temporally aligned relative to each other using the phase of the F1 modulation of thalamic excitation recorded in their postsynaptic L4 target neurons in response to gratings drifting in the preferred and non-preferred direction. Bottom: The average of the phase aligned thalamic excitation recorded in the postsynaptic L4 cortical neurons from the same 8 pairs.

These data imply that for a direction selective L4 cortical neuron, the temporal distribution of activity of the population of its thalamic inputs depends on the order in which the spatial phases of a grating are presented. For a grating drifting in the preferred direction, they should fire together in phase to produce the observed large amplitude fluctuations of thalamic excitation in the postsynaptic neuron. In the opposite direction their firing should be more homogeneously distributed in time. We used our direction selective compound neuron to determine the behavior of the presynaptic population of thalamic neurons to gratings drifting in the preferred and opposite direction (Fig. 4c). The responses of the six L4 neurons contributing to the compound neuron, and of their eight presynaptic thalamic input neurons, were temporally aligned using the phase of the F1 modulation of thalamic excitation (see Methods). Importantly, there was no correlation between the DSI of the firing of individual thalamic neurons and the DSI of their postsynaptic L4 cortical target neuron (Extended Data Fig. 7). In the preferred direction the eight thalamic neurons fired in phase to produce a strongly F1 modulated population activity (Fig. 4c). In the opposite direction the same thalamic neurons fired in a more distributed manner resulting in a much less pronounced F1 modulation of the population activity. Thus, our data demonstrate that the convergence of transient and sustained thalamic neurons responding to distinct spatial phases produces a spatiotemporal offset that confers direction preference to their target L4 cortical neuron.

A simple model in which two thalamic neurons, a transient and a sustained, preferring distinct spatial phases of the stimulus, converge on the same cortical neuron, captures the essence of the above observations (Fig. 5). Because of their transient and the sustained firing, the two thalamic neurons generate a fast and a slow decaying EPSC onto the cortical neuron, respectively, whose amplitude depends on the spatial phase of the stimulus. Because the sustained and the transient neuron prefer distinct stimulus phases the decay of the compound EPSC in the cortical neuron changes with the phase of the stimulus. The direction preference and the DSI of the cortical neuron depend on the relative offset of the spatial phases of the two converging thalamic neurons (Fig. 5).

**Figure 5.**
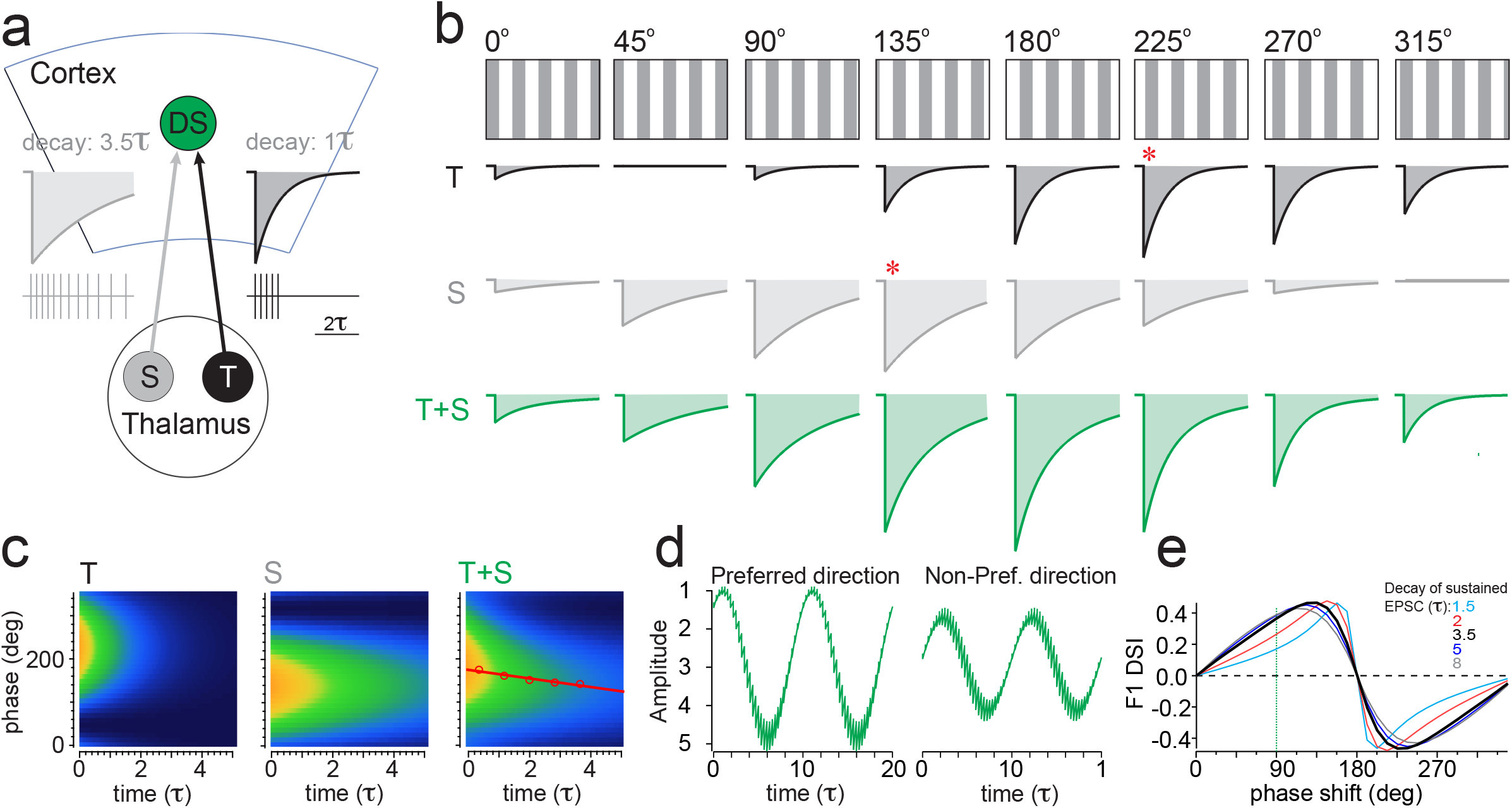
A simple model of direction selectivity. a. Schematic of the model: A cortical neuron (DS) receives two excitatory inputs, one that generates currents with a fast decay (1 tau; dark gray) and the other that generates currents with a slower decay (3.5 tau; light gray). The slow and the fast decays of the currents reflect the sustained (S) and the transient (T) firing of the thalamic input neurons in response to a visual stimulus, respectively. The receptive fields of the two thalamic neurons are offset in space relative to each other. b. The excitatory currents mediated by the T and S neurons and their sum (T+S; green) in response to static gratings presented at distinct spatial phases. Because the receptive fields of the two thalamic neurons are offset (by 90 degrees in phase space for this particular example) the currents that each generate in the cortical neurons peak (red asterisks) in response to distinct phases of the static grating. As a consequence of this offset, the decay of the compound EPSC (T+S) depends on the spatial phase of the stimulus. c. The spatiotemporal receptive field of the currents generated by the T and the S neuron and by their sum (T+S). The spatiotemporal receptive field of the sum but not of the individual components is tilted. d. Response of the cortical neurons to gratings drifting in either direction. The cycle period is 10 tau. e. The DSI of the cortical neuron is plotted against the spatial phase difference (phase shift) of the T and S neurons for the example above (black) and for different decays of the EPSC generated by the sustained thalamic input neuron (colored lines). The vertical green line marks the 90 degrees shift of the example above.

Finally, our 23 paired recordings allow us to estimate the number of thalamic neurons that contribute to the response of a L4 neuron to drifting gratings (Fig. 6). We convolved the spike train of each thalamic unit in response to drifting gratings with its uEPSC (Fig. 6a), integrated the resulting current and compared the obtained charge with the total thalamic charge received by its postsynaptic L4 neuron in response to drifting gratings (Fig. 6b). This metric hence combines the size of the uEPSC relative to the total thalamic excitation and the firing rate of the presynaptic thalamic unit. A large contribution of an individual thalamic unit (unitary contribution) may be due to a large uEPSC and/or a high firing rate. The unitary contribution averaged 1.25 ± 1.51% (n = 23 pairs). Randomly sampling the pool of 23 unitary contributions indicate that on average 80±10 thalamic neurons contribute to the excitation of a L4 neuron (Fig. 6b; see Extended Data Fig. 8 for an estimate of the range). The unitary contribution was the same no matter whether the stimulus drifted in the preferred or opposite direction (1.25 ± 1.54% preferred; 1.25 ± 1.51% opposite direction; p=0.52 Wilcoxon signed rank test), consistent with the lack of direction selectivity of the thalamic charge.

**Figure 6.**
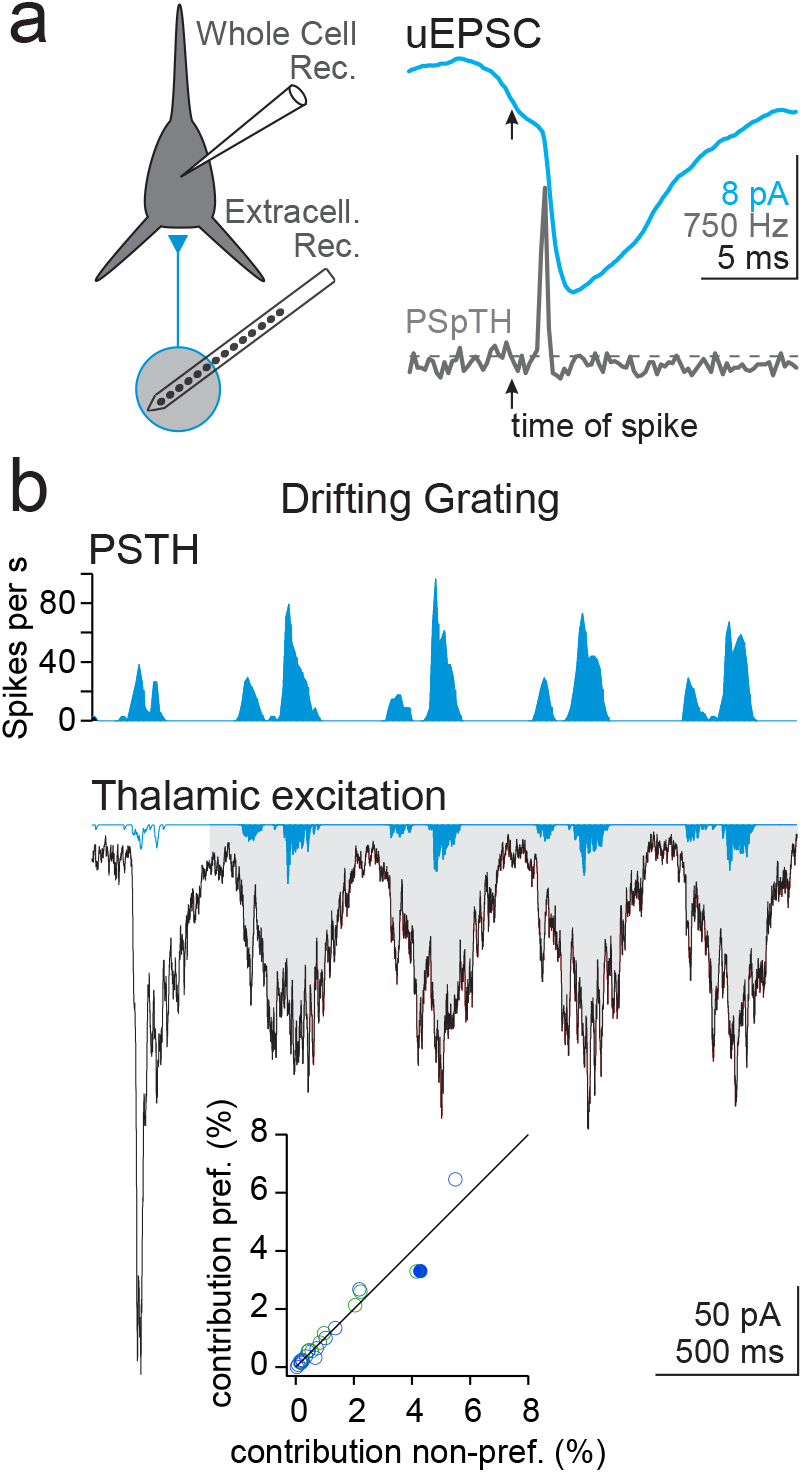
Contribution of individual thalamic neurons to thalamic excitation in visual cortex. a. Left: Schematic of experimental configuration. Right: A unitary excitatory postsynaptic current (uEPSC) from a presynaptic thalamic unit recorded in a L4 cortical neuron while silencing cortex. The arrows indicate the time of the spike. The gray trace is the peri spike time histogram (PSpTH) of events recorded in the L4 neuron (see methods and Extended Data Fig. 5). b. Top: Peristimulus time histogram (PSTH) of this thalamic unit in response to a drifting grating. Bottom: Black trace: Thalamic excitatory current recorded in the L4 neuron in response to drifting grating. Blue trace: Unitary thalamic excitation in response to drifting grating computed by convolving the uEPSC with the PSTH. Shaded areas are the time integral of thalamic excitation. This thalamic neuron contributed ~4% (unitary contribution) of total thalamic excitation. Inset: comparison of unitary contributions computed for the preferred and the non-preferred direction for all thalamo-cortical pairs (n=23). Green circles: from cortical neurons with DSI of thalamic excitation >0.3 (n = 8). Blue circles from cortical neurons with DSI<0.3 (n = 15). Filled blue circle is the example above.

## Discussion

These results show that the initial seed for the generation of direction preference in L4 neurons of V1 originates from the combination of thalamic inputs with distinct spatiotemporal response properties. Thus, the direction of motion is extracted from the visual scene at the earliest stage of cortical processing by integrating thalamic inputs that convey information about distinct spatial locations with distinct time courses. While these results reveal the mechanism for the primordial bias in direction selectivity in visual cortex other intracortical mechanisms may further shape and amplify this bias. These mechanisms may include a sharpening by anisotropic intracortical inhibition or excitation ^10,15,16,18–21^, as well as a sharpening and amplification by cortical recurrent excitatory circuits ^15,31,34^, by the intrinsic excitable properties of the neuronal membrane ^11^ and the active properties of dendrites ^38^. In contrast, the fact that the isolated thalamic charge is, on average, the same irrespective of the direction of the stimulus (Fig. 1g,h), that equalizing the thalamic charge for the two directions of the stimulus has a minor impact on the DSI of F1Thal (Extended Data Fig. 2) and that the direction preference of a cortical neuron does not correlate with the direction bias of its thalamic inputs (Extended Data Fig. 7) shows that direction selectivity in earlier, pre-cortical stages of visual processing, e.g. the retina, play a minor role in the emergence of direction selectivity in L4 neurons. However, we cannot exclude the possibility that anesthesia may have reduced the impact of other mechanisms in the emergence of directional selectivity. We optogenetically excite inhibitory neurons to silence visual cortex and this perturbation may even extend beyond the borders of V1. Most known excitatory input to L4 in V1 originate from either within V1 or from the dLGN and thus we believe that most visually evoked excitation isolated during cortical silencing originates from the dLGN. Other sources of visually evoked excitation from yet uncharacterized cortical or sub-cortical sources, whose response to visual stimuli does not depend on V1 may have, however, contributed to the recorded excitation.

The two classical models for the detection of directional motion by the nervous system, the Reichardt ^12^ and the Barlow and Levick ^13^ models are based on spatiotemporal offsets in excitation or in inhibition relative to excitation, respectively. Both types of offsets have been proposed as a mechanism for the generation of direction selectivity in visual cortex ^10,15,16,18–20^. Our results indicate that the spatiotemporal offset of excitatory inputs relative to each other, originally proposed in the Reichardt model, more closely captures the mechanism for the emergence of direction preference in mouse L4 cortical neurons.

Differences in the time-course of the visual response of thalamic neurons, categorized as transient and sustained^39^ as well as lagged and non-lagged ^23,40^, have been reported in several species including cats^39,23,40^, monkeys ^41^ and mice ^37^, where they were originally proposed to contribute to direction selectivity^23,24^. Thus, the mechanism identified in the present study may be generalizable to other species. Some of the dLGN neurons categorized here as sustained had a late onset reminiscent of lagged cells ^23,40^ (Extended Data Fig. 4). While our model (Fig. 5) only considers transient and sustained dLGN neurons with the same onset, late onset dLGN neurons could further enhance the direction selectivity of the cortical neuron.

The distribution of uEPSC amplitudes in our dataset is skewed towards smaller values and large amplitude EPSCs were likely undersampled. Furthermore, submillisecond synchrony between thalamic units may have led us to misclassify some thalamic units as presynaptic to the recorded L4 neuron when they in fact only fired some of their action potential in synchrony with an actual presynaptic unit. These two factors may have led to an underestimate of the average uEPSC amplitude and hence an overestimate of the number of units contributing to the visual response of a L4 neuron (Extended Data Fig. 8). Nevertheless, compared to previous thalamo-cortical paired recordings our estimate of uEPSP amplitude is very similar to that reported in rodent somatosensory cortex *in vivo* ^35^ and *in vitro* ^42^, and somewhat larger than in cat visual cortex *in vivo* ^36^. Furthermore the convergence of thalamic neurons onto L4 cortical neurons estimated here (~80) resembles estimates in somatosensory cortex based on paired recordings *in vivo* ^35^ suggesting a common thalamo-cortical convergence across distinct sensory systems.

In conclusion, in the mammalian nervous system direction selectivity is generated *de novo* in at least two stages of visual processing, the retina and the cortex. While, based on studies in primates and carnivores, it was originally believed that retinal direction selectivity was not transmitted to the geniculo-cortical pathway but instead exclusively conveyed to other subcortical structures, recent evidence in rodents indicates that dLGN neurons can inherit direction selectivity from the retina^27,37,43^ and that they project to the cortex ^28,29^. However, abolishing direction selectivity in the retina does not eliminate direction selectivity in cortex ^30^ consistent with its *de novo* emergence in cortex described here. Whether direction selectivity computed in the cortex through the mechanisms illustrated here is combined, at some stage of cortical processing, with direction selectivity inherited from the retina or whether these two channels stay separated remains to be established.

## Methods

Experiments were performed in accordance with the regulations of the Institutional Animal Care and Use Committee of the University of California, San Diego and the Administrative Panel on Laboratory Animal Care at Stanford University.

### Mice

Mice were heterozygous male or female offspring of PV-Cre ^44^, JAX #008069) or VGat-ChR2 ^45^, JAX#014548) transgenic mice crossed with ICR white wildtype animals. Data were obtained from 41 VGat-ChR2 mice, and 7 PV-Cre mice. The number of cells and mice for each experiment are detailed below and in the main text.

Drifting grating current clamp and voltage clamp recordings (Fig. 1a-g): 52 cells, 31 VGat-ChR2 mice. Drifting grating voltage clamp recordings (Fig. 1h, Extended Data Fig. 1c-d): 61 cells in 36 VGat-ChR2 mice; 5 cells in 5 PV-Cre mice.

Drifting grating/static grating voltage clamp recordings (Fig. 2) 48 cells in 29 Vgat-ChR2 mice; 5 cells in 5 PV-Cre mice.

Drifting grating/static grating V1 voltage clamp/LGN silicon probe recordings (Fig. 3-5, Extended Data Fig. 3-4): 40 dual recordings in 24 Vgat-ChR2 mice.

Cortical extracellular recording (Extended Data Fig. 1a): 2 PV-Cre mice; 5 Vgat-ChR2 mice.

Static grating LGN silicon probe recording (Extended Data Fig. 4): 24 recordings in 24 Vgat-ChR2 mice.

### Solutions

Artificial Cerebrospinal Fluid (ACSF): 140 mM NaCl, 5 mM KCl, 10 mM D-glucose, 10 mM HEPES, 2 mM CaCl2, 2 mM MgSO4, pH 7.4.

Potassium-based intracellular solution: 135 mM potassium gluconate, 8 mM NaCl, 10 mM HEPES, 4 mM Mg-ATP, 0.3 mM Na-GTP, 0.3 mM EGTA, pH 7.4.

Cesium-based intracellular solution: 125 mM cesium methanesulphonate, 8 mM NaCl, 10 mM HEPES, 4 mM Mg-ATP, 0.3 mM Na-GTP, 0.3 mM EGTA, 2 mM QX-314, pH 7.4ChR2 expression in V1 inhibitory interneurons

### ChR2 expression in V1 inhibitory interneurons

AAV1-Flex-ChR2-tdTom (University of Pennsylvania Vector Core) was injected in the visual cortex of neonatal (P0-1) PV-Cre mice to achieve ChR2 expression in PV+ inhibitory interneurons (PV-ChR2 mice) as previously described ^31^ VGAT-ChR2 mice express ChR2 in cortical inhibitory interneuron populations therefore no viral injections were required.

### Surgery for recording experiments

Adult mice (5-12 weeks) were anesthetized with a combination of urethane (1.5 g/kg, IP), chlorprothixene (2-4 mg/kg, IP), and light isoflurane (0.5% in O2). A drop of silicon oil was applied to the eyes. The scalp was removed, the skull cleaned and a metal head-fixation bar was affixed to the skull using dental acrylic. A 1-2 mm diameter craniotomy was performed over V1 in one hemisphere (1 mm anterior of the lambdoid suture, 2.5 mm lateral to the midline). In simultaneous V1 and LGN recording experiments, a second narrow elongated craniotomy was performed (spanning 2-3 mm posterior of bregma, 3.4 mm lateral to the midline) for insertion of the silicon probe array into the LGN. In both craniotomies the dura was removed. Recording began shortly after completion of surgery.

### V1 whole cell recording

Whole cell recordings were made using the blind patch technique ^46^ at a depth of 300-550 μm corresponding to layer 4. Patch pipettes with 4-6 MΩ resistance were pulled from borosilicate glass and filled with intracellular solution. Potassium-based intracellular solution was used in 54 cells (52 recorded in both current clamp and voltage clamp, 2 recorded in voltage clamp only). 12 cells were recorded with cesium-based intracellular solution (voltage clamp only). Prior to insertion of patch pipettes, a drop of low-melting point agarose (1.5% in ACSF) was applied to the brain surface to reduce movement. Pipettes were rapidly inserted into the visual cortex while applying high positive pressure (2.5 psi). Pressure was reduced to 0.5 PSI upon reaching a depth of 200-300 μm and the pipette was advanced in 2 μm steps while monitoring the pipette resistance. When a sudden increase in resistance was encountered, positive pressure was released and light suction was applied to achieve a gigaohm seal. Brief pulses of suction were applied to break the seal and achieve whole-cell configuration. Series resistance was 20-50 MΩ. Signals were amplified (Multiclamp 700B, Molecular Devices) and digitized at 10 kHz (DigiData, Molecular Devices) or 31.25 kHz (PCIe-6259, National Instruments). Membrane potential and spiking responses were recorded in current clamp configuration. Excitatory currents were recorded in voltage clamp at -70 mV near the reversal potential of inhibition.

### LGN unit recording

Prior to initiating V1 whole-cell recordings, a 4-shank 32 channel silicon probe (Buzsaki32, NeuroNexus) was inserted into the LGN craniotomy (coordinates listed above) with the shanks distributed along the anterior/posterior axis. The probe was inserted at a 55 degree angle above horizontal in the coronal plane such that it advanced along the lateral to medial and dorsal to ventral directions. The probe was advanced in 5 μm steps until robust visual responses were observed in the multiunit activity (2.5-2.8 mm distance). The shanks typically spanned the anterior/posterior extent of the dLGN from 2-3 mm posterior of bregma. Retinotopy as assessed with coarse receptive field mapping of multiunit activity was consistent with Piscopo et al, 2013. Signals were amplified (Model 4000, AM Systems) and digitized at 31.25 kHz (PCIe-6259, National Instruments). For technical reasons, the top-most site on each shank was not recorded. The probe was allowed to settle for 30-60 minutes before collecting data. At the end of the experiment, the mouse was sacrificed under deep anesthesia and the brain was fixed in 4% paraformaldehyde. The tissue was sectioned for post-hoc verification of the recording sites.

### Retinotopic alignment of V1 and LGN recordings

To improve the chances of recording from synaptically connected LGN and V1 neurons, coarse receptive field maps of the LGN multiunit activity were generated and the corresponding retinotopic region of V1 was identified by mapping the receptive field of the LFP signal at various locations in the V1 craniotomy. The number of V1 locations that were sampled before finding the corresponding cortical location was approximately 1-5 although this was not precisely documented during the experiments. Subsequent V1 whole-cell recordings were targeted to this region of V1. V1 LFP was recorded using a patch pipette filled with ACSF inserted to a depth of 200-400 μm.

### Cortical silencing

Visual cortex was silenced by illuminating the V1 craniotomy with a 1mm fiber optic coupled to a blue LED (470 nm; 20 mW total output, Doric) positioned several mm above the craniotomy or through the objective (20x) of a fluorescence microscope with a blue LED (470 nm, 2.3 mW total output, Thorlabs) coupled to the excitation port. The LED turned on 650 ms prior to the onset of a visual stimulus trial and lasted throughout the duration of the visual stimulus. Trials with cortical silencing were interleaved with trials without illumination in which cortical activity was intact. To validate the effectiveness of cortical silencing, spiking responses of V1 neurons in response to drifting gratings were recorded using loose-patch or silicon probes (Buzsaki32 or A1×32-Edge-5mm-20-177, NeuroNexus) during the silencing protocol. Cortical silencing in both PV-ChR2 and VGat-ChR2 mice suppressed nearly all spiking of non-narrow spiking neurons in V1 (~99% suppression, Extended Data Fig 1b). Some of the experiments illustrated in Extended Data Fig 1b appear in previous publications and were also performed by one of the authors (PV-ChR2: all loose patch recordings are from Fig.1b in ^31^;VGat-ChR2: 64 out of 138 units are from Supplemental Fig. 7g-h in ^47^).

### Visual stimuli

Visual stimuli were presented on an LCD monitor (75 cd/m^2^ mean luminance, gamma corrected) to the eye contralateral to the hemisphere in which recordings were performed.

### Drifting gratings

Drifting gratings stimuli were full-field, full contrast drifting bar gratings (0.04 cyc/deg spatial frequency, 2Hz temporal frequency). For determining the preferred orientation, 12 different drifting grating stimuli were presented consisting of 6 evenly-spaced orientations (30 degree increment) drifting in one of two opposite directions along the axis perpendicular to the grating bars. Drifting gratings were presented for 2.3 s and were preceded and followed by a mean luminance gray screen. Presentation of a single grating was considered one visual stimulus trial (see cortical silencing methods). In 52/66 recordings, the preferred orientation and direction of the cell were determined in current clamp using the full set of 12 orientations/directions followed by recording in the voltage clamp configuration using drifting grating stimuli restricted to the 2 opposite directions at the preferred orientation. In these cells the orientation of the stimulus may not exactly match the preferred orientation of thalamic excitation. In the remaining 14 cells the recording was started in the voltage clamp configuration and the preferred orientation was determined in voltage clamp during cortical silencing by presenting the full set of 12 orientations/directions. The orientation and/or direction of the grating stimuli were presented in random order.

### Static gratings

Static grating stimuli were full-field, full contrast bar gratings (0.04 cyc/deg spatial frequency) of the preferred orientation of the cortical neuron. 16 evenly-spaced spatial phases were presented (22.5 deg increment equal to 1/16 of a cycle). A series of 5 static gratings of randomly chosen spatial phase were presented sequentially. Each static grating was presented for 0.25 s and followed by a mean luminance gray screen for 0.25 s. Each 5-grating sequence was considered one visual stimulation trial (see cortical silencing methods) and was preceded and followed by a mean luminance gray screen.

### Direction selectivity index

Direction selectivity index (DSI) for responses to two drifting grating stimuli moving in opposite directions was calculated as:

> (RespPref - RespNull)/RespPrefDir

Where RespPref and RespNull are the size of the responses to the directions that gave the larger and smaller responses, respectively. This limits the range of DSI from 0-1. The responses used to calculate the DSI were the spike rate, the F1 amplitude modulation of Vm, the F1 amplitude modulation of the thalamic excitation and the thalamic excitatory charge (see Drifting grating analysis below). When the DSI was compared for two different response parameters within the same cell e.g. DSI of the F1 modulation of thalamic excitation versus the DSI of the F1 modulation of Vm (Fig. 1g middle), the DSI was defined relative to the preferred direction of one of the parameters, the reference parameter. If the preferred direction of the two parameters were different, the DSI of the non-reference parameter was multiplied by - 1. In the example above the reference parameter was Vm. Hence, the DSI of the reference parameter is always positive (range 0 to +1) but the DSI of the non-reference parameter can be negative or positive (range -1 to +1) with negative and positive values indicating that the two parameters prefer opposite or the same direction, respectively. The absolute value of the DSI indicates the degree of selectivity.

### Drifting grating analysis

Drifting grating responses were evaluated in a time window from 0.3 s after stimulus onset to the end of the stimulus (2 s total or 4 complete cycles). Spikes in current clamp recordings were detected by identifying time points where the membrane potential exceeded -15 mV. The time of a spike was defined as the time of the peak depolarization in a 1.5 ms window following each threshold crossing. Prior to calculating the DSI of F1 amplitude for subthreshold membrane potential responses (Fig. 1g), all spikes were removed from current clamp recordings by replacing the time window from 1 ms before to 2 ms after the spike time with linear interpolation. In voltage clamp recordings, the holding current in a 0.4 s window prior to the onset of visual stimulus was subtracted for each trial, computed from the bottom 5th percentile of the distribution of current values, which should include the periods with the least amount of spontaneous excitatory activity. The DSI based on spikes was calculated from the average spike rate during the response window. F1 amplitude, for both Vm and thalamic excitation, was derived from the amplitude of a sinusoidal fit to the cycle average of drifting grating responses. The period of the cycle average was 0.5 s, matching the temporal frequency of the drifting grating. The excitatory charge was calculated from the integral of drifting grating excitatory current responses.

### Static grating analysis

Analysis was restricted to responses collected under cortical silencing conditions (thalamic excitation). The thalamic excitation in response to each spatial phase of the static grating were averaged together. The average value of the current from 0-24ms after stimulus onset was subtracted. This time window was prior to any observable visual response.

### Static grating summation

The algebraic sum of thalamic EPSCs evoked by each of the 16 phases of the static gratings was computed for each cortical neuron. Before computing the sum, the 16 EPSCs were staggered in time relative to each other in proportion to the phase offset between the phase of the static grating used to evoke each of the EPSCs and a fixed reference phase. The absolute time separation was 31.25 ms for every 22.5 degrees of phase offset such that a full cycle (16*31.25 ms) corresponds to 500 ms, i.e. a full cycle of a 2Hz drifting grating. We computed the sum for positive and negative time separations (+ or - 31.25ms) corresponding to the temporal sequence of phases of a grating drifting in one or the opposite direction. The DSI and F1 amplitudes of the resulting algebraic summations were analyzed as described above.

### Static grating spatiotemporal analysis

Spatiotemporal analysis was restricted to cortical neurons whose summed static grating responses predicted the preferred direction in response to drifting gratings (DSI summed static *relative* to drifting > - 0.1). The spatial phases were ordered such that the relationship between the sequence of spatial phases and the preferred direction in response to drifting gratings was the same for each neuron. Specifically, the preferred direction is in the upward direction for static grating heatmaps (Fig. 2f and 4a,b).

### Early/Late spatiotemporal analysis

Early and late thalamic excitation was defined as the excitatory charge from 30-110 ms and 110-230 ms after the onset of the static grating stimulus, respectively. The preferred spatial phase of early and late thalamic excitation was calculated from the vector average across all spatial phases of early or late thalamic excitation, respectively. For population average of early and late thalamic excitation (Fig. 2e right), the early and late thalamic excitation were upsampled in the spatial phase dimension by 4x and shifted along the spatial phase axis so that the preferred spatial phase of early thalamic excitation occurred at a spatial phase of 180 degrees.

### Spatiotemporal slope analysis

The preferred spatial phase was calculated for 5 time bins (40ms width starting from 30ms after stimulus onset). A linear fit to these 5 preferred spatial phase values was performed to calculate the spatiotemporal slope. For population averages of static grating heatmaps (Fig. 2f right, Fig. 4b), each cortical neuron’s static grating responses were normalized by the peak response across all spatial phases, upsampled in the spatial phase dimension by 4x and shifted along the spatial phase axis so that the preferred spatial phase of the earliest time bin (30-70ms) occurred at a spatial phase of 180 degrees.

### Thalamic unit spike sorting

The UltraMegaSort ^48,49^ spike sorting software was used to detect, cluster, and assign spike waveforms into single units as previously described ^50^. Spike waveforms on the 7 recorded sites of an individual shank were clustered using a *k*-means algorithm followed by manual assignment of clusters with distinct waveform profiles into single units.

### Identification of monosynaptically-connected thalamocortical pairs

For each simultaneously recorded thalamic unit and cortical neuron, a brief segment of the excitatory current recorded in the cortical neuron during drifting grating visual stimulation with cortical silencing around the time of each thalamic spike was extracted (25 ms before and after the spike). We looked for the presence of a fast inward (negative) deflection in thalamic excitation occurring 1-4 ms after the thalamic spike. We first calculated the first derivative of thalamic excitation of the spike-triggered sweeps (dtThExc). The spike-triggered average of dtThExc was computed and z-scored. Thalamic units containing a z-score peak exceeding -5 in the time window of 1-4 ms after the thalamic spike were considered candidate presynaptic units. To ensure that the negative peak in the spike-triggered average of dtThExc was due to an actual increase in the probability of fast inward events occurring in the 1-4 window following the spike, we detected the occurrence of such events in the spike-triggered dtThExc sweeps. Events were detected as decreasing threshold crossings of dtThExc where the threshold was set as - 1*(standard deviation of all spike-triggered dtThExc sweeps for each thalamic unit). A peri-spike time histogram (PSpTH) was assembled from these event detections (0.2 ms bin size) and z-scored. Candidate presynaptic units with a z-score peak exceeding 3.5 in the time window 1-4 ms after the thalamic spike were considered monosynaptically connected to the cortical neuron. Differentiation was performed on data that was resampled at 10 kHz using consecutive samples. Following differentiation, sweeps were smoothed (0.3 ms running average) to generate dtThExc. Z-scores of the spike-triggered average of dtThExc and PSpTH were computed by first subtracting a smoothed version (3 ms running median to remove fluctuations slower than several ms) and then z-scoring using the average and standard deviation of time points 0-5 ms prior to spike.

Latency of monosynaptic response was defined as the time of the peak in the z-scored PSpTH relative to the spike. Jitter was defined as the half-width at half-max of this peak. The unitary EPSC (uEPSC) was derived from the spike-triggered thalamic excitation of each monosynaptically-connected thalamic unit. While the identified monosynaptic connections had sub-millisecond jitter, the uEPSC may ride on top of a slower envelope of thalamic excitation driven by the response of other synaptically connected inputs to the visual stimulus. This component was estimated by shifting the trial number of thalamic responses by 1 trial relative to that of thalamic excitation for each direction of the drifting grating and computing a spike-triggered average based on the trial-shifted data^35^. The trial-shifted spike-triggered average was subtracted from the uEPSC (shift-subtracted uEPSC) for amplitude and contribution (see “Thalamic unit contribution” below) analyses. The baseline from 0-1 ms after the thalamic spike was also subtracted. Unsubtracted uEPSCs are shown in the main figures. Shift-subtracted uEPSCs are shown in Extended Data Fig. 3. The amplitude of monosynaptic connections was defined as the most negative (inward) value of the shift-subtracted uEPSC from 0-3 ms after the onset of monosynaptic response (i.e., the time of thalamic spike + latency for each pair) minus the average value in a 0.2 ms window just prior to the onset. Amplitude of uEPSP and spontaneous uEPSC were determined in the same manner using the shift-subtracted spike-triggered average of Vm during drifting gratings under control conditions or the spike-triggered average of thalamic excitation during the 500ms prior to visual stimulation during cortical silencing, respectively. Spontaneous uEPSCs were only characterized for pairs with at least 30 spontaneous spikes. One thalamic unit that passed the criteria for monosynaptic connection did not exhibit a clear spontaneous uEPSC in the simultaneously recorded cortical neuron despite firing a sufficient number of spontaneous spikes and hence was not considered to be monosynaptically connected. In the 4 experiments in which multiple presynaptic thalamic neurons were recorded, the spike sorting of monosynaptically connected thalamic units was verified using KiloSort software ^51^. For one of these experiments, this yielded an additional thalamic unit which passed the criteria for monosynaptic connection.

In all subsequent connected pair analyses, each pair was treated independently without regard for whether or not the postsynaptic neuron was common to other connected pairs. Thus the postsynaptic responses of a cortical neuron may be represented multiple times if more than one pre-synaptic thalamic unit to that cortical neuron was identified (e.g. the pairs in Fig. 3a).

### Connected pair static grating analysis

Out of 23 pairs, 20 of the pre-synaptic thalamic units had spiking responses to static gratings. For each pre-synaptic thalamic unit, peri-stimulus time histograms (PSTH) of the spiking were constructed for responses (10 ms binning, upsampled to 10 kHz, smoothed by 20 ms running average) to each spatial phase of the static grating under cortical silencing conditions.

### Duration of thalamic excitation response to static gratings

The duration was defined as the time point after stimulus onset at which the excitatory *charge* reached 90% of its maximum value. In Figure 3, analyses were restricted to those static grating responses in connected pairs in which both the pre-synaptic LGN firing and the postsynaptic thalamic excitation were at least 10% of that elicited by the spatial phase that gave the largest response. Each response was peak normalized.

To test statistical significance of the average pairwise Pearsons correlation between thalamic excitation (EPSC) and thalamic spiking (PSTH) spatial phase responses, the PSTH responses across all pairs and spatial phases were shuffled relative to their corresponding EPSC response so that each EPSC response was reassigned to a random PSTH response. For each shuffle, the average pairwise correlation was calculated, and this was procedure was repeated for 10,000 shuffles. The average pairwise correlation of the real data was compared to the distribution of shuffled average pairwise correlations. None of the shuffled average pairwise correlations exceeded that of the real data. Use of the Spearman correlation produced similar results.

### Compound cortical neuron: static grating

For each pair, the static grating heatmaps of the presynaptic PSTH and the postsynaptic thalamic excitation were shifted along the spatial phase axis so that the preferred spatial phase of the earliest time bin (30-70ms) of thalamic excitation occurred at a spatial phase of 180 degrees and the sequence of spatial phases was ordered so that the preferred direction of thalamic excitation in response to drifting gratings was in the upward direction. Heatmaps were normalized by the peak response across all spatial phases, upsampled in the spatial phase dimension by 4x, and averaged across all pairs.

### Compound cortical neuron: drifting grating

Cycle-average responses to drifting gratings of the preferred and non-preferred direction of thalamic excitation of all pairs were averaged together. Before averaging, the EPSC and PSTH cycle-average for each pair was shifted in time by the same amount so that the F1 peak of thalamic excitation occurred at 250 ms and peak normalized.

### Thalamic unit contribution

For each pair, the excitatory current contributed by the thalamic unit was calculated by convolving its shift-subtracted uEPSC from 0-15 ms after the thalamic spike with its spike train during drifting gratings and averaging across trials. The shift-subtracted uEPSC was truncated to the time point at which it returned to baseline if this occurred before 15 ms. The contributed charge was the integral of the trial-averaged convolution across the stimulus duration. Dividing the contributed charge by the thalamic excitatory charge evoked by the same drifting grating stimulus resulted in the fractional contribution of the thalamic unit.

### Statistical analysis

Statistical analyses were done in IgorPro. No statistical methods were used to pre-determine sample sizes, but our sample sizes were similar to those reported in previous publications in the field. All data are presented as mean ± s.d. Normality of the data were not tested and nonparametric two-sided Wilcoxon rank-sum or Wilcoxon signed-rank tests were used for unpaired or paired tests, respectively. Fraction of cells with matching direction preference was compared to a chance value of 0.5 using two-tailed binomial test. Experiments and analysis were not blinded.

### Code availability

Custom code used in this study are available from the corresponding author upon reasonable request.

### Data availability

The datasets generated during and/or analyzed during the current study are available from the corresponding author upon reasonable request.

## Acknowledgments

We thank J. Evora for help with genotyping and mouse husbandry, S. Hestrin for allowing us to perform some of the experiments in his lab, R. Beltramo for helping with extracellular recordings, J.S. Isaacson and B.L. Bloodgood for comments on the manuscript and the members of the Scanziani and Isaacson laboratories for helpful discussions of this project. This project was supported by the Gatsby Charitable Foundation and the Howard Hughes Medical Institute.

## Author contributions

A.D.L. and M.S. designed the study. A.D.L. conducted all experiments and analysis. A.D.L. and M.S. wrote the paper.

**Extended Data Figure 1.**
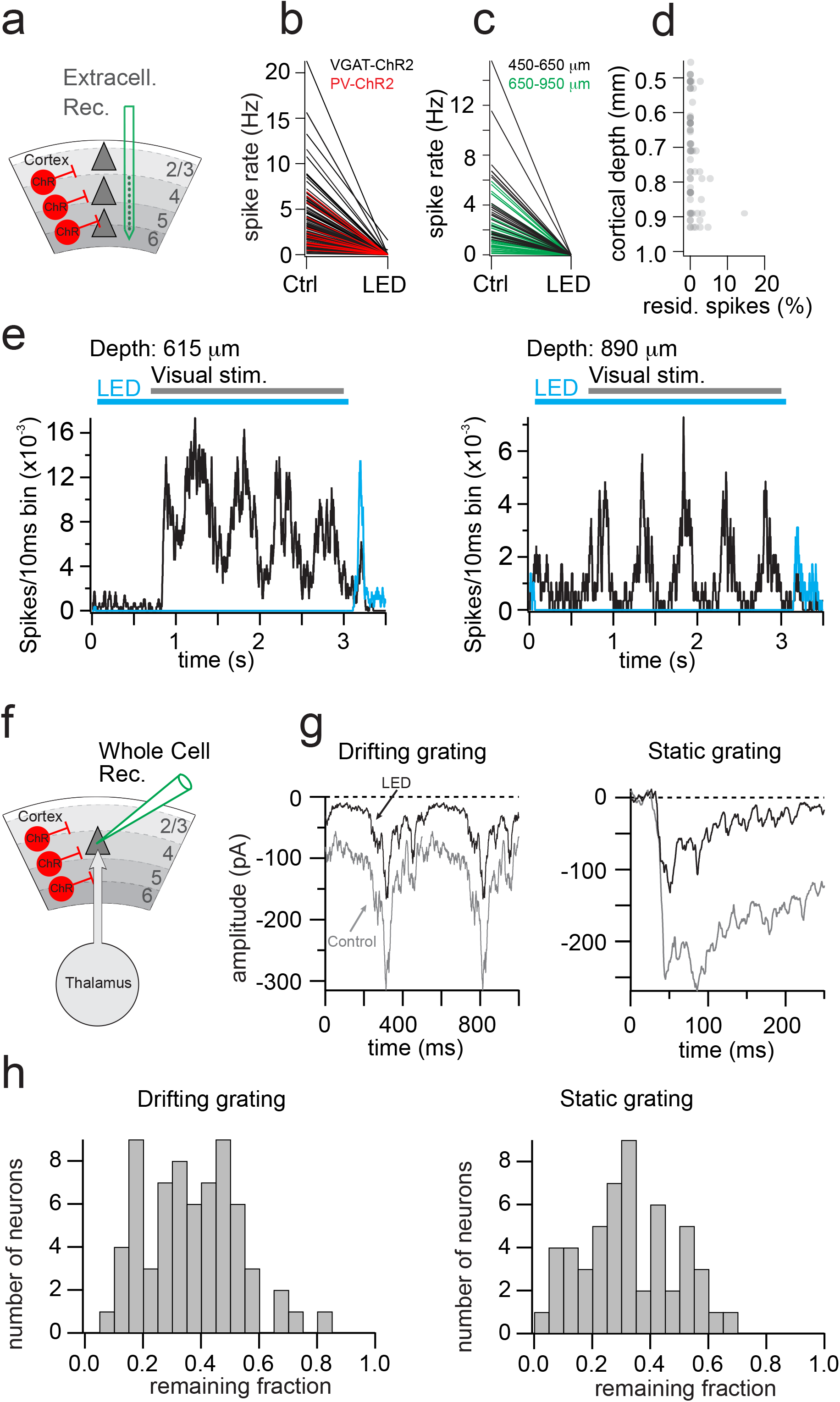
Cortical Silencing. a. Experimental configuration: Channelrhodopsin (ChR2) is expressed in cortical inhibitory neurons to suppress neuronal activity upon illumination with a blue LED while performing extracellular recordings. b. Visually evoked activity (full field drifting gratings) from units isolated throughout the cortical depth is suppressed upon LED illumination. Black lines: ChR2 was expressed into all GABAergic neurons (vGat-ChR2 mouse line; 138 units recorded with silicon probes; 25 units recorded above 500 μm from the cortical surface (98.9 ± 2.7% silencing) and 113 recorded below 500 μm (99.4 ± 2% silencing); 5 mice). Red lines: ChR2 was conditionally expressed in parvalbumin expressing neurons through viral injection into the visual cortex of the PV-Cre mouse line (13 loose patch recordings in layer 4; 2 mice; 100% silencing). c. As in (b) but specifically for units recorded between 450-650 μm depth (black; 99.8+/− 0.6% silencing; n = 26; 3 mice) and 650-950 μm (green; putative layer 6; 99.8+/− 2.4% silencing; n = 48; 3 mice) from the cortical surface. These units are a subset of the units from vGat-ChR2 mice illustrated in (b) where the exact recording depth could be estimated. All units from vGat-ChR2 mouse line. d. Percent visually evoked spikes remaining during LED illumination across cortical depths deeper than 450 um. Same units as in (c) e. Peristimulus time histogram of two units located at 615 μm (left) and 890 μm depth (right) in response to drifting gratings under control conditions (black) and during cortical silencing (blue). The duration of the visual stimulus and of the LED illumination is illustrated by the horizontal bars. f. Experimental configuration: as in (a) but whole cell recordings from layer 4 neurons instead of extracellular recordings. g. Whole cell voltage clamp recording (V holding: -70 mV) of a layer 4 neuron (same neuron as in Fig. 2a). Cycle average in response to drifting gratings (left; two identical cycles are shown for clarity) and to static gratings (right; average of 10 traces). Gray: Control conditions. Black: During LED illumination to isolate the thalamic component of excitation. h. Distribution of residual excitatory charge upon LED illumination for drifting gratings (66 recordings similar to (g) left) and static gratings presented at the preferred spatial phase (53 recordings similar to (g) right).

**Extended Data Figure 2.**
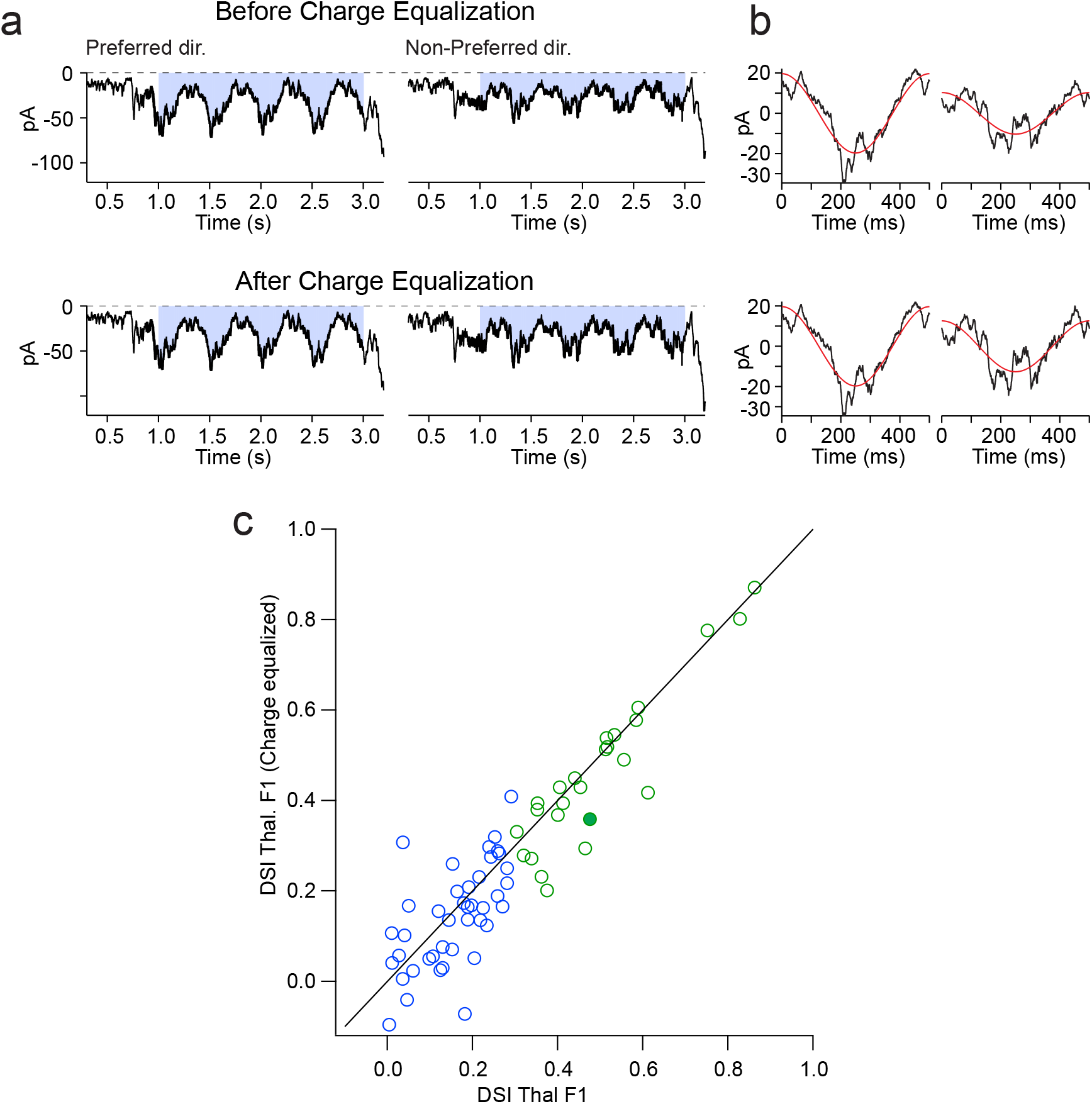
The contribution of the directional preference of the thalamic charge to the direction selectivity index of ThalF1. In those neurons in which the preferred direction of the thalamic charge matched that of VmF1 (positive DSI Thal. Charge; Fig. 1g right) the absolute value of the DSI of the thalamic charge (0.087+/−0.053; n=32) was significantly different than the absolute value of the DSI of the thalamic charge in those neurons in which the preferred direction of the thalamic charge did not match that of VmF1 (negative DSI Thal. Charge; Fig. 1g right; 0.048+/−0.037 (n=20); p= 0.003; t-test). To determine the impact of this slight bias in thalamic charge on DSIThalF1 we have equalized the thalamic charge evoked by gratings drifting in both directions. a. Example recording from a cortical neuron where the charge of thalamic excitation is larger in the preferred as compared to the non-preferred direction. Top: thalamic excitation as recorded (non equalized) in response to a grating drifting in the preferred (left) and non-preferred (right) direction. Bottom: same as top but after scaling the response to the non-preferred direction such that the charge is the same in either direction (charge equalization). b. Cycle average of thalamic excitation with superimposed sinusoidal fit (red). Top: as recorded; Bottom: after charge equalization. After charge equalization the direction preference is maintained but, for this particular example, the DSI is reduced (see filled datapoint in (c)). c. Scatter plot for all recordings (Green: DSI> 0.3; Blue DSI<0.3). The filled data point is the example above. The equalization leads to only a very small change in the DSI of the thalamic F1 amplitude modulation (All neurons: DSI before equalization: 0.28 +/− 0.20; DSI after equalization: 0.26 +/− 0.21 (p=0.034; paired t-test; n=66); Subset of neurons where DSIThalF1 > 0.3 (green): DSI before equalization: 0.49 +/− 0.15; DSI after equalization: 0.46 +/− 0.17 (p=0.022; paired t-test; n=25); Subset of neurons where DSIThalF1 < 0.3 (blue): DSI before equalization: 0.16 +/− 0.09; DSI after equalization: 0.14 +/− 0.11(p=0.300; paired t-test; n=41).

**Extended Data Figure 3:**
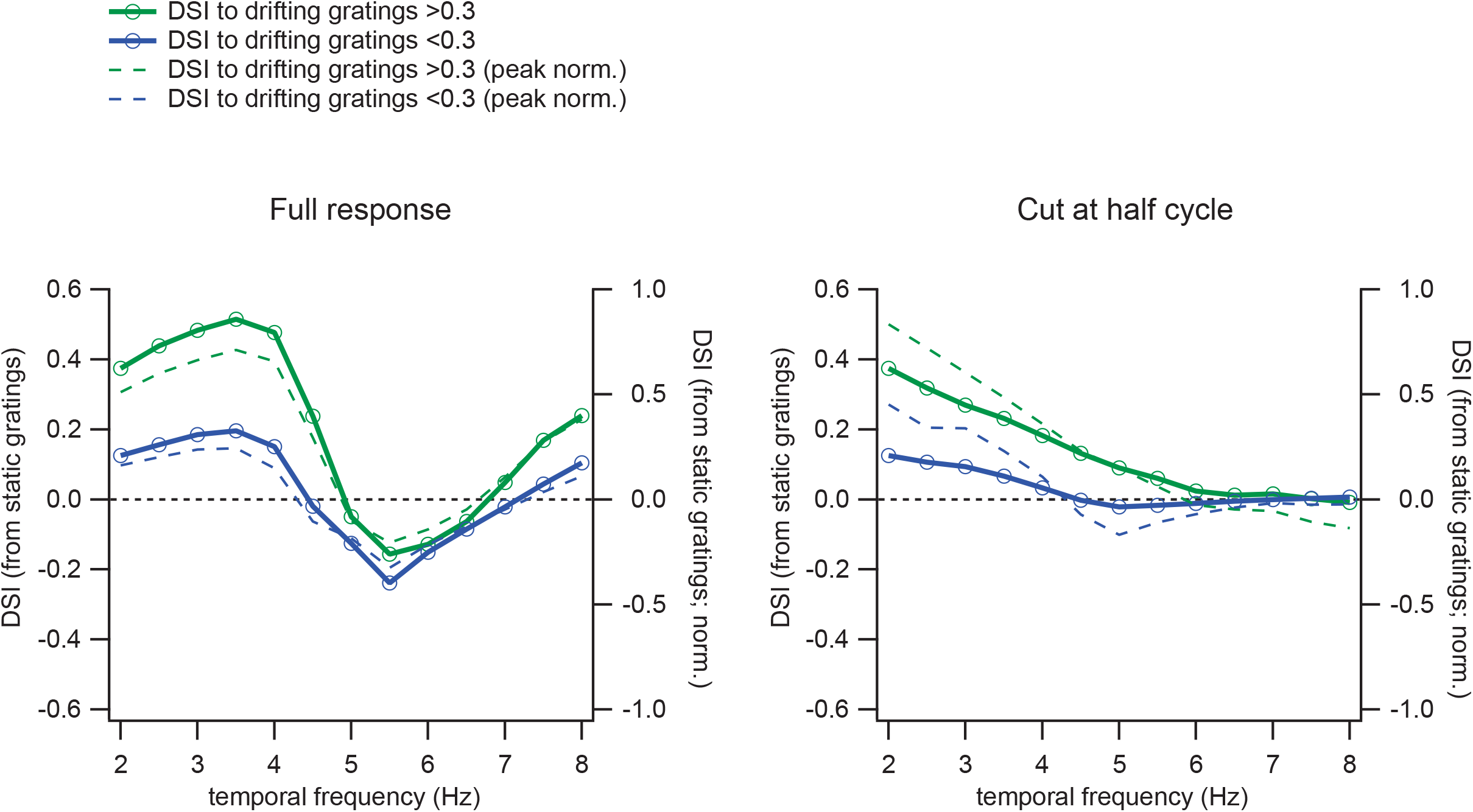
Predicting the DSI for Various Temporal Frequencies. Direction selectivity index (DSI) predicted from the response to static gratings. The amplitude of the F1 modulation was determined from the algebraic sum of the thalamic EPSCs evoked by each of the 16 phases of the static grating. The thalamic EPSCs were staggered in time to mimic different temporal frequencies of a drifting grating (e.g. at 4Hz a cycle lasts 250 ms and hence the response to each one of the phases is staggered by 15.6 ms (250/16 ms) relative to the preceding one). The DSI was computed by comparing the F1 modulation of the sum in which EPSCs were ordered according to the spatial phase sequence simulating the motion of the grating in one direction against the sum simulating motion in the opposite direction. Green and blue traces, average of all cells whose DSI ThalF1 to drifting gratings was larger or smaller than 0.3, respectively. Dotted traces: the computed DSI was normalized to the peak for each cell; right ordinate. *Left panel:* the full 250 ms response to static grating was used to compute the DSI at each temporal frequency. Note the reversal of direction preference at higher temporal frequencies. *Right panel:* Only the initial x milliseconds of the response to static gratings were used to compute the DSI, x being the half period of the temporal frequency to be computed (e.g. for 4 Hz, x = 125 ms). The rationale for this approach is that the interactions between excitatory inputs that are relevant for the emergence of DS likely occur within a half cycle.

**Extended Data Figure 4.**
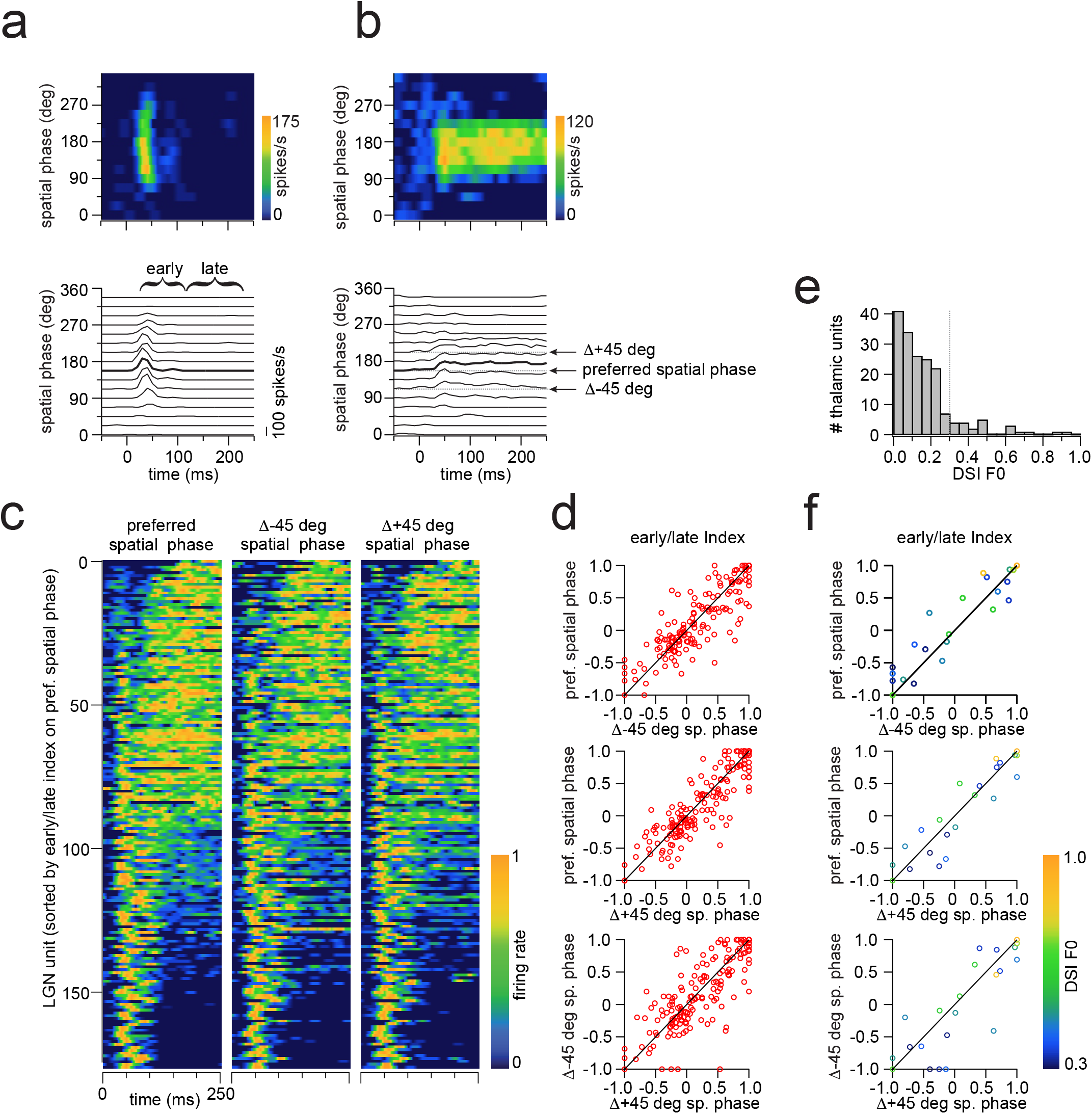
The response dynamics of dLGN units to static gratings is similar across phases. a. Example dLGN unit with transient response to static gratings. Top: Spatiotemporal receptive field. Bottom: Peri-stimulus time histograms (PSTHs) in response to each phase of the static grating used to construct the spatiotemporal receptive field illustrated above. The PSTH at the preferred phase is highlighted by a thicker trace. The preferred phase is defined as the phase closest to the vector average of the response at each phase. The brackets show the time windows over which the early and late firing rates were averaged (Re and Rl, respectively) in order to compute the early/late index [(Re-Rl)/(Re+Rl)]. This unit has an early/late index of 1 for static gratings presented at the preferred phase and of 0.88 and 1 for gratings presented at phases of ± 45 degrees from the preferred phase. b. As in (a) but for an example dLGN unit with sustained response to static grating. The arrows illustrate the preferred spatial phase and the phases separated by ±45 degrees. This unit has an early/late index of -0.3 for static gratings presented at the preferred phase and of -0.08 and 0.22 for gratings presented at phases of ± 45 degrees from the preferred phase. c. Heat-maps of responses to static gratings for 177 thalamic units (24 mice). Left: Each row is the amplitude of the PSTH of one of the dLGN units in response to the preferred phase of the static grating. The units are ordered according to their early/late index in response to the preferred phase. Middle: Same as left but in response to a static grating whose phase is 45 degrees below the preferred phase. The order of the units has not been changed; i.e. it is the same as on the left. Right: Same as left but in response to a static grating whose phase is 45 degrees above the preferred phase. The order is the same as on the left. Note that transient and sustained units maintain their characteristic firing dynamics even in response to static gratings presented at phases of ±45 degree from the preferred phase. d. Scatter plots of the early/late index computed in response to static gratings presented at the preferred phase and at phases of ±45 degrees from the preferred. Note that in all plots the data are close to the unity line. e. Distribution of direction selectivity indexes of the firing rates (DSI F0) of dLGN units in panel d. The vertical dotted line is DSI F0 = 0.3. f. As in (d) but specifically for those dLGN units with a DSI F0 larger than 0.3 (n=22 units).

**Extended Data Figure 5.**
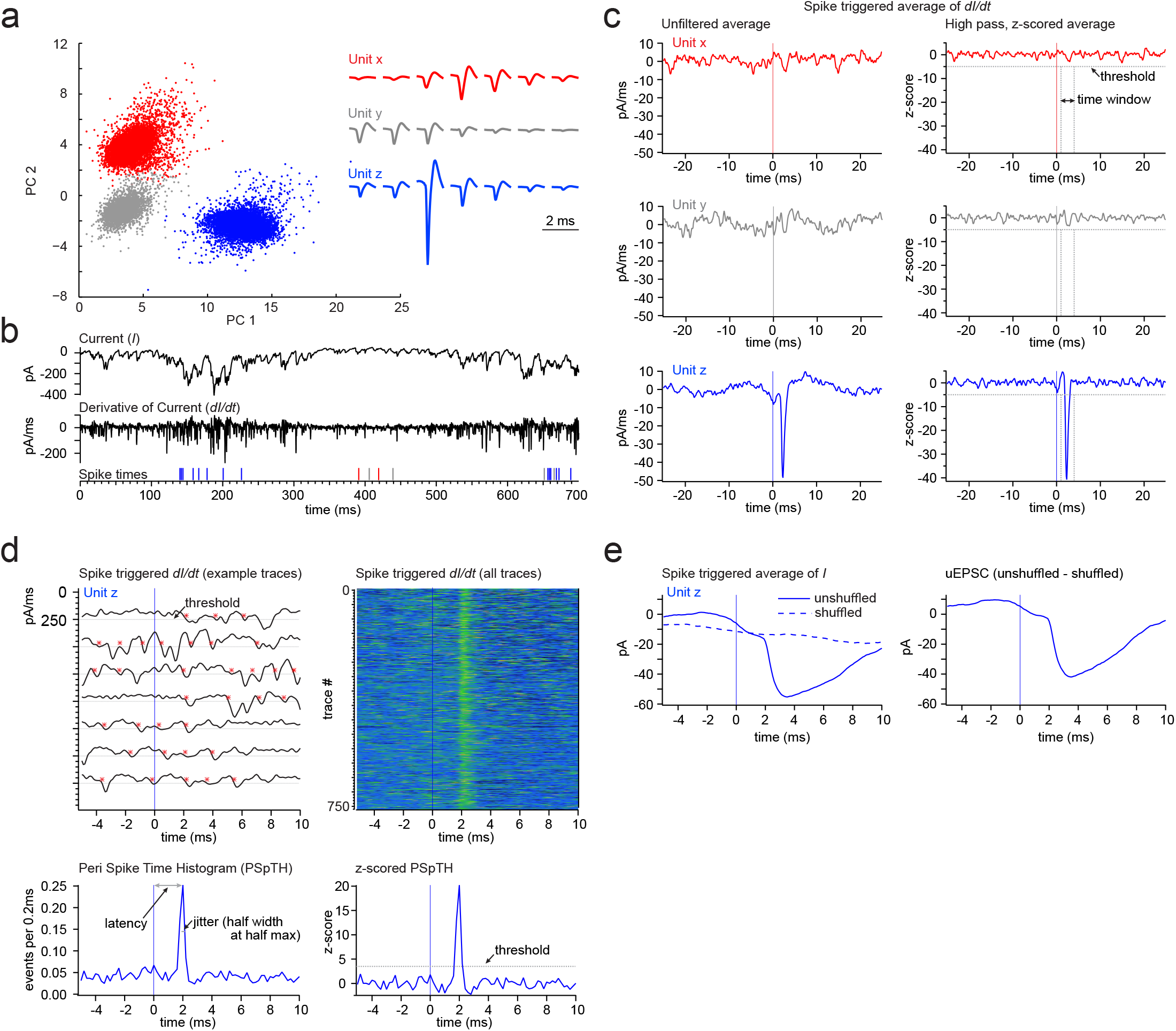
Criteria for identifying thalamocortical pairs. Thalamocortical pairs were identified based on two criteria: Criterion 1 (illustrated in panel c) sets a threshold for the dLGN unit spike triggered average of the time derivative of the current recorded in L4 cortical neurons. Criterion 2 (illustrated in panel d) sets a threshold and a time window for the distribution of events detected in the time derivative of the L4 current around the time of the spike in the dLGN unit. Both criteria have to be satisfied for the dLGN unit and the L4 cortical neuron to be considered a pair. a. Isolation of units in the dLGN. Right: First two principal components illustrating three separable clusters attributed to three independent dLGN units (units x, y and z in red, gray and blue, respectively). Left: Electrophysiological recording illustrating the average spike shape recorded from 7 electrodes for the three dLGN units. b. Differentiation of the current recorded in L4 neurons. Same experiment as in (a). Top trace: The current recorded in the whole cell configuration from a L4 cortical neuron (holding voltage:-70 mV) in response to the presentation of a drifting grating (single trial). Middle trace: The temporal derivative of the above current *(dI/dt).* Lower panel: the times at which each one of the three dLGN units from (a) (x: red; y: gray; z: blue) fired during the same trial. c. Criterion 1. Left panels: spike triggered average of *dI/dt* of the current recorded in the L4 neuron for the three dLGN units illustrated in (a). Time 0 denotes the time of the spike. Right panels, same spike triggered averages shown on the left after low pass filtering and z-scoring (see methods). Note that only unit z (blue) crosses the 5z threshold. d. Criterion 2. Top left: Seven individual time derivatives of currents *(dI/dt)* recorded in the L4 neuron (same as in (b)) aligned relative to seven spikes recorded in unit z (time 0 denotes the time of the spike). Each asterisk shows an event crossing the threshold of -36pA/ms. Top right: same as left but represented as a heat-map of the amplitude of *dI/dt* for 761 traces (the heatmap color scale ranges from +50 pA/ms to -200 pA/ms). This heat-map clearly illustrates an increase in event probability around 2 ms following the spike in unit z. Bottom left: the Peri Spike Time Histogram (PSpTH) for the events detected in the 761 traces illustrated above. The peak of the PSpTH is used to determine the latency (i.e. the time interval between the spike recorded in the dLGN unit and the occurrence of a postsynaptic response detected in the L4 cortical neuron). The half width at half max is used to determine the jitter of that response (in this example the latency is: 2 ms and the jitter is 188 microseconds). Bottom right: Same as left but z-scored. The PSpTH must cross a threshold of 3.5z within 1 -4 ms after the spike in the dLGN unit for the dLGN unit to be considered synaptically connected to the L4 neuron. e. Left: Unit z spike triggered average of the response recorded in the same L4 neuron as in (a). Continuous blue line: unshuffled trials. Dotted line: shuffled trials (see methods). Right the difference between the shuffled and unshuffled trials is used to isolate the unitary EPSC (uEPSC) between unit z and the recorded L4 neuron.

**Extended Data Figure 6.**
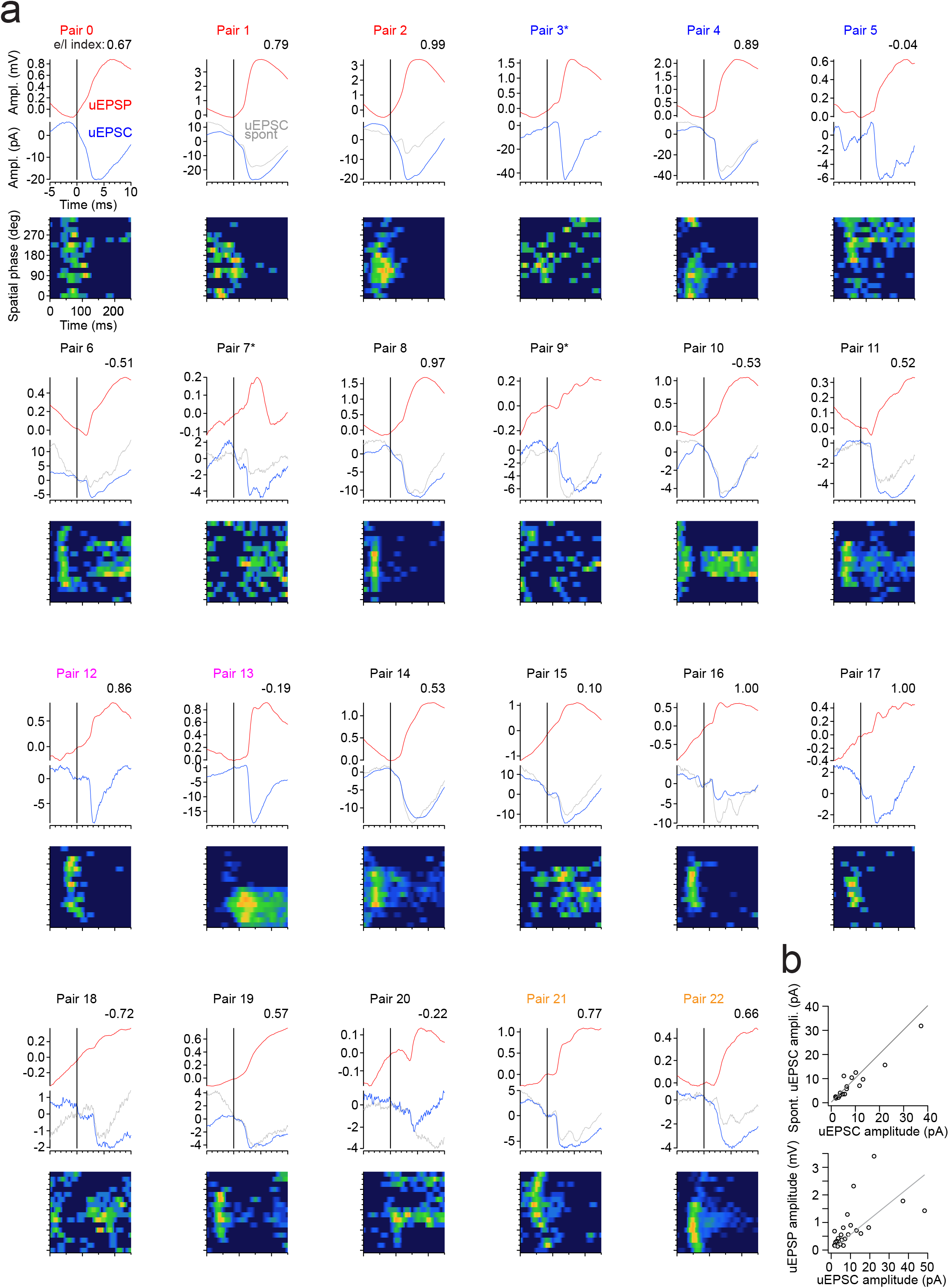
Unitary EPSCs, EPSPs, and spatiotemporal receptive fields of presynaptic thalamic units. a. Each panel shows the shift-subtracted unitary EPSP (uEPSP; top red, see methods), the shift-subtracted unitary EPSC (uEPSC; middle blue, see methods) recorded during visual stimulation and the spatiotemporal receptive field (heat-map; bottom) of one of the 23 thalamic units connected to a layer 4 neuron. For some pairs the uEPSC recorded during spontaneous activity (gray, see methods) is also shown. uEPSCs are recorded during cortical silencing; uEPSPs are recorded under control conditions. The vertical line at time 0 marks the time of the peak of the extracellularly recorded action potential in the presynaptic thalamic unit. Pair numbers of the same color were recorded in the same postsynaptic layer 4 cortical neuron. The heat-map shows the spatiotemporal receptive field of the thalamic unit in response to static gratings. Each spatiotemporal receptive field is centered (157.5 degrees) on the preferred spatial phase (defined as the phase that produced the most spikes) of its unit except for converging pairs which are aligned to the average preferred phase of the converging units. The response of pairs marked by an asterisks were not included in the analysis of static gratings because of their poor response to those stimuli. The early/late (e/l) index (see Extended Data Fig. 4) for the preferred phase of the presynaptic thalamic unit is given for each pair on the top right except for pairs with an asterisk. uEPSCs (blue) are the average of 49 - 970 spike triggered traces. uEPSPs (red) are the average of 101 - 1496 spike triggered traces. Spontaneous uEPSCs (gray) are the average of 30 - 412 spike triggered traces. Units of pairs 10, 11, 12, 13, 14, 18, 21, and 22 are the eight presynaptic units to the compound neuron in Figure 4 and correspond to the units numbered in Figure 4 as 6, 7, 1, 2, 8, 5, 4, and 3, respectively. b. Top panel: Correlation between the amplitude of the visually evoked uEPSC (blue in (a)) and the spontaneously occurring uEPSC (gray in (a); r=0.95 p=5.9e-9 n=17) for those pairs in which both could be recorded. The gray line is unity. Bottom panel: Correlation between the amplitude of the visually evoked uEPSC (blue in (a)) and the uEPSP (red in (a); r=0.59 p=0.0028) for all pairs. The gray line is linear fit to the data with a slope of 0.056 mV/pA.

**Extended Data Figure 7.**
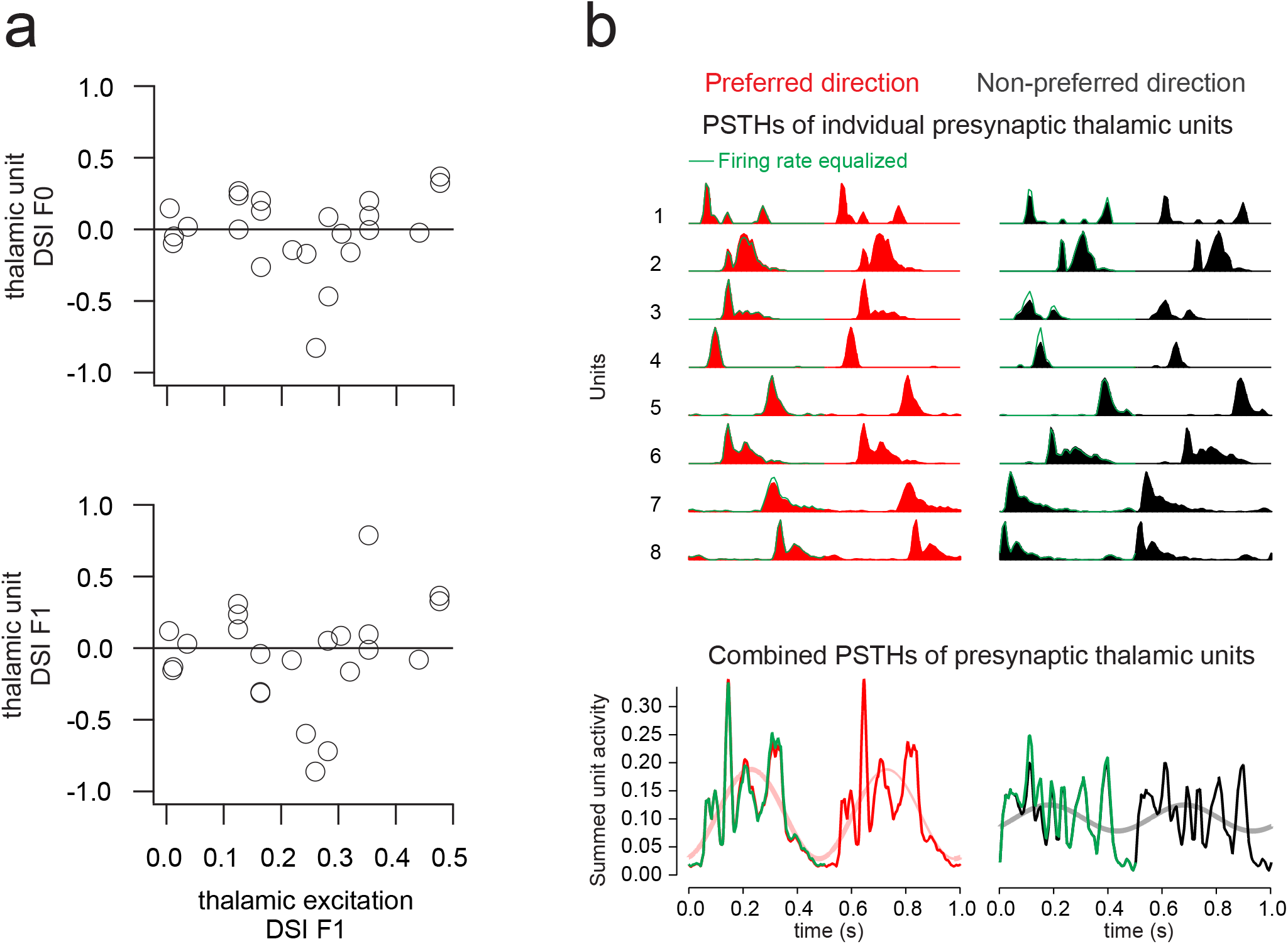
The direction selectivity of presynaptic thalamic units does not contribute to the direction selectivity of layer 4 cortical neurons. a. Top: The DSI of the average firing rate of each presynaptic thalamic unit (thalamic unit DSI F0) is plotted against the DSI of the F1 modulation of thalamic excitation recorded in the postsynaptic L4 cortical neuron (thalamic excitation DSI F1; r=0.09, p=0.68; 12/23 matching preferred direction, p=1.00, binomial test; n = 23 pairs). Bottom: The DSI of the F1 modulation of the firing rate of each presynaptic thalamic unit (thalamic unit DSI F1) is plotted against the DSI of the F1 modulation of thalamic excitation recorded in the postsynaptic L4 cortical neuron (r=0.16, p=0.46; 11/23 matching preferred direction, p=1.00, binomial test; same 23 pairs as left). Note that the DSI of thalamic units does not predict the DSI of the F1 of thalamic excitation. b. Top: The PSTHs of each of the 8 units contributing to the compound neuron (from Fig. 4). The units were temporally aligned relative to each other using the phase of the F1 modulation of thalamic excitation recorded in their postsynaptic L4 target neurons in response to gratings drifting in the preferred (red) and non-preferred direction (black). Two identical cycles are shown for clarity. The equalized PSTHs (i.e. the PSTHs that were scaled such that the firing rate of the thalamic unit is the same in either direction) are shown in green. Only the first cycle is equalized to facilitate comparison. Bottom: Summed PSTHs of the 8 presynaptic thalamic units (pink and gray lines are sinusoidal fits; from Fig. 4). The green traces are the summed activity of the equalized PSTHs. Note the similarity between the control and the equalized combined activity.

**Extended Data Figure 8.**
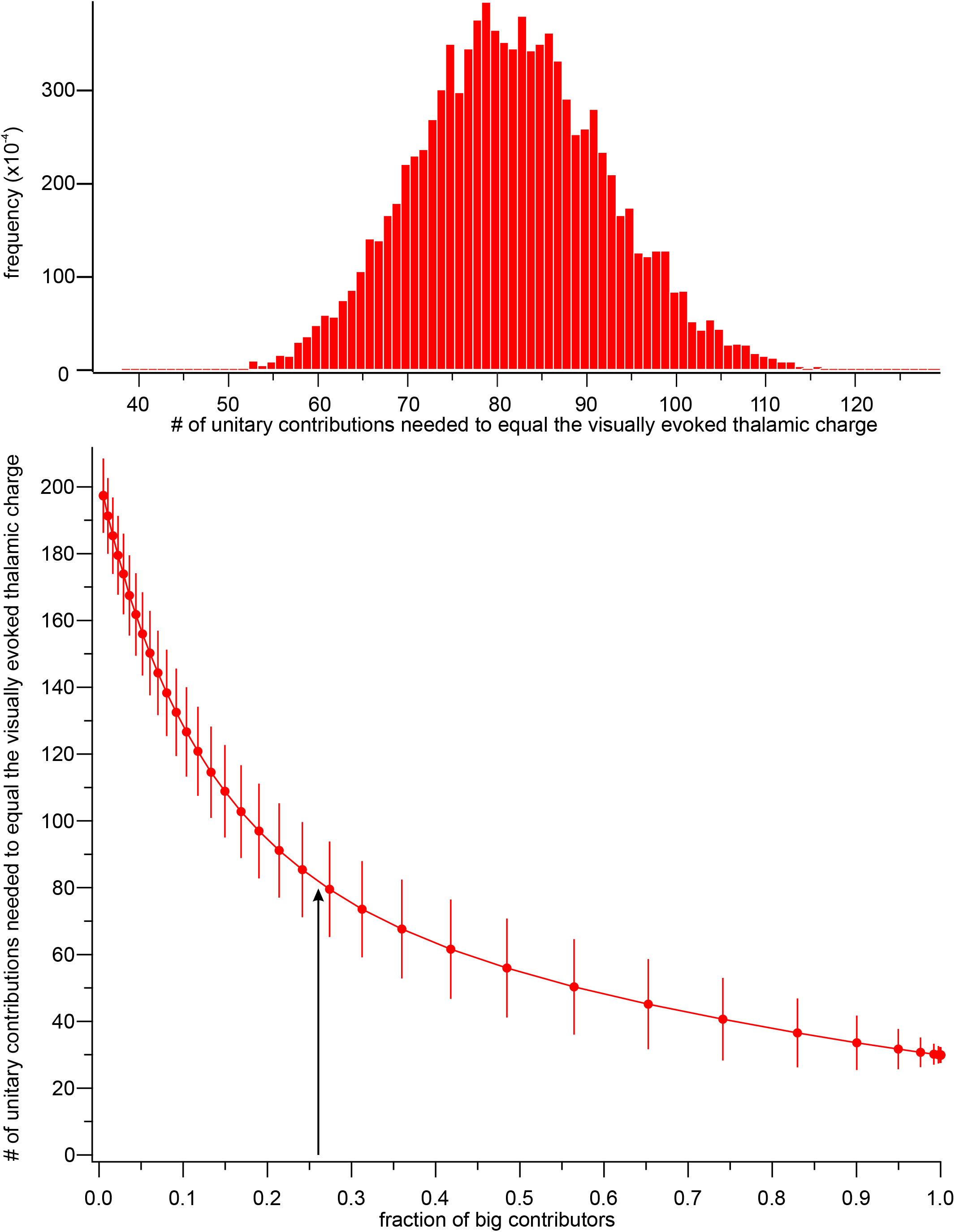
Number of thalamic units contributing to the visually evoked response of a L4 cortical neuron. *Top panel:* Distribution of the number of unitary contributions of dLGN neurons necessary to equal the total thalamic charge recorded in a L4 cortical neuron in response to a grating drifting in the preferred direction during cortical silencing. The distribution is obtained by randomly sampling with replacement the individual contributions from each of the 23 pairs, 10,000 times. In each iteration, unitary contributions were sampled until their sum reached 100%. To compute the unitary contribution of a dLGN unit we first convolved the spike train of the unit in response to a drifting grating with the uEPSC that that unit evoked in the postsynaptic L4 cortical neuron, integrated the resulting current in time and normalized the obtained charge by the total charge recorded in the postsynaptic cortical neuron in response to the drifting grating during cortical silencing. Unitary contributions are expressed in percent of the total charge. On average 80.9 +/− 10.7 dLGN units (average +/− std) contribute to the visually evoked thalamic current in a L4 cortical neuron. *Bottom panel:* Number of unitary contributions of dLGN neurons necessary to equal the total thalamic charge as a function of the fraction of “big contributors”. Because the units that contribute a large fraction of the total charge (big contributors) may have been under-sampled (as a consequence of a skewed distribution) we have arbitrarily increased their fraction in the pool of unitary contributions and determined the average number of unitary contributions necessary to equal the total charge, as above. Big contributors are those dLGN units that contribute to more than 2% of the total charge. They represent 26% of all unitary contributions in our data set of 23 pairs (6 pairs; arrow). Increasing the fraction of big contributors (x-axis) progressively reduces the average number of dLGN units necessary to equal the total thalamic charge evoked in response to visual stimulation (y-axis). Each data point is the average +/− sem.

